# Co-targeting Metabolic Neighbours Constraints Bacterial Adaptive Evolution

**DOI:** 10.64898/2026.06.04.729935

**Authors:** Dwipanjan Sanyal, Yun-Ti Chen, Carl Ho, Arthur McClelland, Sourav Chowdhury, Eugene I. Shakhnovich

## Abstract

Antibiotic resistance often arises through genetic changes that reduce drug susceptibility, with mutation representing one of the primary mechanisms. Here, we report a dual targeting strategy that restricts these escape routes by simultaneously inhibiting two metabolic neighbours in folate biosynthesis pathway. To establish a general framework for limiting adaptive escape in bacteria, we screened the folate pathway to identify enzymes with shared conserved structural features. This analysis identified dihydroneopterin aldolase (folB) and dihydroneopterin triphosphate 2′-epimerase (folX) as optimal dual targets due to their shared pocket architecture and their roles early in the folate pathway. We then screened a large chemical space of FDA-approved small molecules to identify candidates predicted to bind both folB and folX, followed by experimental validation. We identified Arbutin showing the strongest growth inhibition in *E. coli* and emerging as the potent dual-target inhibitor, validated by metabolic rescue assays that confirmed direct inhibition of both folB and folX. Experimental evolution further revealed that populations exposed to Arbutin exhibit constrained adaptive trajectories and minimal increases in tolerated drug concentration, consistent with the evolutionary bottleneck imposed by dual-enzyme inhibition. Together, these results establish a framework for developing evolution-resistant antibacterials by targeting structurally convergent enzymes within metabolic pathways.

## Introduction

Antibiotic resistance remains one of the most urgent biomedical challenges of the modern era(Chinemerem Nwobodo, Ugwu et al. 2022, Salam, Al-Amin et al. 2023). Despite continued discovery of new antimicrobial therapeutics, bacterial populations continue to evolve adaptive solutions that challenge therapeutic efficacy(Davies and Davies 2010, Theuretzbacher and Piddock 2019, Chinemerem Nwobodo, Ugwu et al. 2022). A major limitation of current drug-development pipelines is the short evolutionary lifespan of single protein targeting compounds, which often select for rapid mutational escape along highly accessible adaptive routes(Davis, Plowright et al. 2017). As a result, resistance can arise through one or a few substitutions that preserve enzyme function while reducing drug sensitivity. Together, these challenges motivate antimicrobial strategies that prioritize evolutionary constraint over maximal biochemical potency, with the aim of restricting the mutational routes accessible under drug selection.

Metabolic pathways offer a promising context for such strategies(Murima, McKinney et al. 2014). Many essential pathways comprise structurally or functionally related enzymes that operate sequentially to generate key metabolites(Walsh 2003, Murima, McKinney et al. 2014, Brown and Wright 2016). Perturbation of these enzymes can impose strong physiological effects by constraining flux through downstream biosynthetic networks(Belenky, Jonathan et al. 2015). Notably, enzymes within the same pathway often exhibit structural convergence despite limited sequence similarity, reflecting shared substrate chemistry, conserved reaction mechanisms, and evolutionary constraints imposed by pathway architecture. Such convergence can result in partially overlapping active-site geometries among enzymes acting on chemically related intermediates. Importantly, structural similarities among enzymes within a pathway presents potential opportunities for multitarget inhibition, in which a single small molecule targets more than one catalytic site(Hopkins 2008, Zhang, Chowdhury et al. 2021). Such coordinated inhibition can impose stringent selective pressure, particularly when mutations that weaken inhibition of one enzyme fail to compensate for inhibition of the other.

Resistance evolution can be visualized as navigation on a high-dimensional adaptive landscape shaped by accessible mutational steps under selection. For single protein targeting inhibitors, this landscape typically contains many low-cost routes that preserve function while reducing drug sensitivity, enabling rapid escape(Rodrigues and Shakhnovich 2019, Boumahdi and de Sauvage 2020). By contrast, simultaneous perturbation of functionally coupled proteins (which can be viewed as nodes of the metabolic or other network) imposes correlated constraints that reduce the number of viable adaptive trajectories and narrow the accessible region of genotype space(Manhart and Shakhnovich 2018). This forces populations toward rarer, higher-cost solutions and slows evolutionary adaptation of pathogens to stressors.

The folate biosynthesis pathway is a conserved metabolic module in prokaryotes and many lower eukaryotes(Appling 1991). In bacteria, it supplies tetrahydrofolate cofactors required for one-carbon transfer reactions that enable nucleotide biosynthesis, amino-acid metabolism, and methylation(Rossi, Amaretti et al. 2011, Toprak, Veres et al. 2011, Rodrigues, Bershtein et al. 2016). Classical antifolates, including trimethoprim and sulphonamides, target essential nodes within this pathway but are prone to rapid resistance evolution, often via binding-site mutations, gene amplification, or adaptive reorganization of metabolic flux(Sköld 2001). Recent works have shown that resistance dynamics in the folate pathway are shaped by its biochemical architecture, with certain enzymes permissive to mutation and others forming evolutionary bottlenecks that restrict escape (Rodrigues, Bershtein et al. 2016, Wistrand-Yuen, Knopp et al. 2018, Baquero, Martinez et al. 2021). These observations suggest that coordinated inhibition of functionally related folate enzymes could restrict accessible mutational pathways.

Using comparative sequence and structural analyses, we focused on two early folate enzymes, dihydroneopterin aldolase (folB) and dihydroneopterin triphosphate epimerase (folX)(Pribat, Blaby et al. 2010). Although folB and folX share only limited primary-sequence identity, they exhibit notable structural similarity, including conserved active-site geometries that accommodate closely related pterin intermediates. Both enzymes catalyse steps in early folate precursor processing(Haußmann, Rohdich et al. 1998), suggesting that coordinated inhibition could disrupt pathway flux at its entry point. This combination of structural convergence and functional coupling indicates that small molecules capable of engaging both enzymes may exert pathway-level control through dual target engagement(Chowdhury, Zielinski et al. 2023).

To explore this possibility, we applied a pathway-guided drug repurposing strategy to identify small molecules capable of simultaneously targeting both enzymes. This approach identified candidate dual inhibitors, among which Arbutin, a naturally occurring glycosylated hydroquinone derivative, emerged as a potent suppressor of bacterial growth. Biochemical and genetic analyses confirmed that Arbutin directly engages both enzymes and perturbs early folate precursor flux.

The central question that extends beyond target inhibition is whether simultaneous inhibition of folB and folX restricts the evolutionary routes available to resistance. Long-term selection experiments revealed sharply divergent adaptive responses to dual inhibition versus single-target inhibition. Populations evolving under inhibition of either enzyme alone rapidly acquired tolerance, whereas those under dual inhibition adapted slowly and incompletely and maintained sensitivity across the drug gradient. These contrasting outcomes were associated with distinct physiological signatures and fitness costs, indicating that coordinated inhibition of folB and folX imposes a substantially more restrictive adaptive landscape. To resolve the genetic mechanisms underlying these contrasting evolutionary outcomes, we performed whole-genome sequencing of evolved populations. We observed that dual inhibition redirects adaptation toward distinct, higher fitness cost genomic routes compared with single-target inhibition.

Together, these results establish a pathway-guided framework for multitarget inhibitor design and demonstrate how coordinated inhibition of structurally convergent metabolic nodes can suppress evolutionary escape, pointing to a general strategy for developing antimicrobial therapeutics with enhanced evolutionary durability.

## Results

### Overview of a common drug strategy for co-targeting the folate biosynthesis pathway

The concept of multi-targeting in drug discovery has been established as an effective strategy for pathway-level drug identification(Dey, Tergaonkar et al. 2008, Jenwitheesuk, Horst et al. 2008, Hsu, Cheng et al. 2013). Based on the hypothesis that multi-targeting can be used to identify key targets for co-targeting within the folate biosynthesis pathway (Figure 1A), we performed a comprehensive sequence and structural analysis of the proteins in this pathway. We identified folX and folB as not only sharing limited sequence homology (24% sequence identity) but also exhibiting high structural similarity (RMSD: 1.97) (Figure 1B). The function of folB is that it facilitates the transformation of 7,8-dihydroneopterin (DHNP) into 6-hydroxymethyl-7,8-dihydropterin (HP). It can utilize both L-threo-dihydroneopterin and D-erythro-dihydroneopterin as substrates to produce 6-hydroxymethyldihydropterin. Additionally, it can efficiently catalyse the epimerization at carbon 2’ from dihydroneopterin to dihydromonapterin(Ahn, Byun et al. 1997, Haußmann, Rohdich et al. 1998). On the other hand, the function of folX is that it facilitates the rearrangement of carbon 2’ in the side chain of 7,8-dihydroneopterin triphosphate (H_2_NTP) to produce 7,8-dihydromonapterin triphosphate (H_2_MTP). The compound structures of DHNP (substrate of folB) and H_2_NTP (substrate of folX) are similar in that both have guanine. In terms of their biological function, their substrates also display a high degree of similarity (Figure 1C). Therefore, we introduce a scoring function, Common Target folB and folX (ComBX), *S*_ComBX_(*p*), where *p* is the top pose and defined in equation (1). It derived from their shared protein–ligand interaction features, to enable the repurposing of co-targeting drugs for folB and folX (Figure 1D). ComBX quantifies protein-ligand binding compatibility based on conserved interaction patterns shared between the folB and folX active sites.

**Figure 1.**
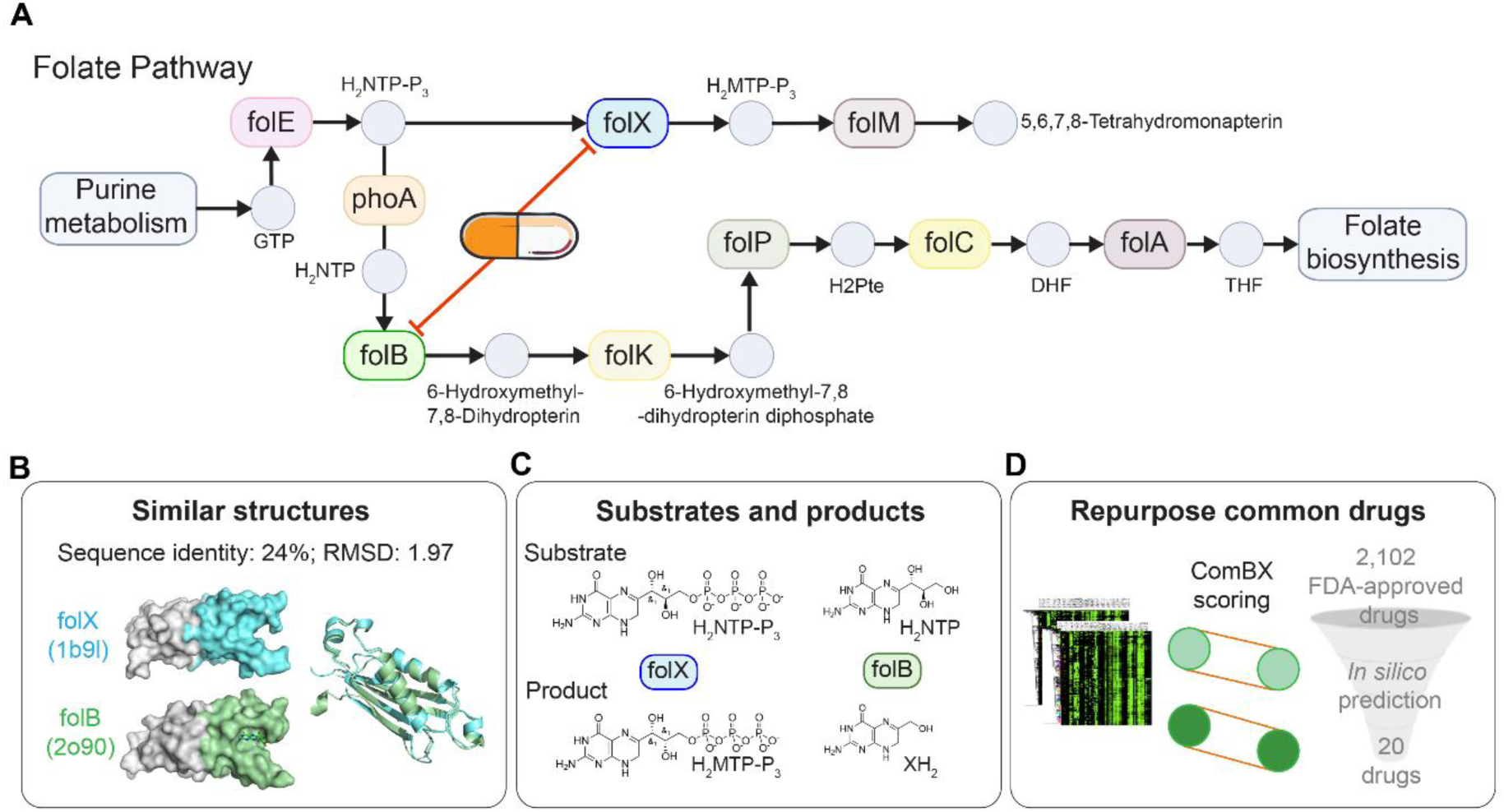
Evolutionary and structural analyses identify two structurally conserved metabolic nodes in the bacterial folate pathway. **(A)** Schematic of the bacterial folate biosynthesis pathway (KEGG pathway: 00790), highlighting key enzymes involved and the idea of a drug for co-targeting. **(B)** Comparison of folB and folX at the sequence and structural levels. Despite low sequence identity (24%), the two enzymes adopt highly similar three-dimensional structures (RMSD = 1.97 Å. **(C)** Chemical reactions catalyzed by folB and folX. folB converts 7,8-dihydroneopterin (H_2_NTP) into 6-hydroxymethyl-7,8-dihydropterin (XH_2_) and glycolaldehyde, while folX converts 7,8-dihydroneopterin 3′-triphosphate (H_2_NTP-P_3_) into 7,8-dihydromonapterin 3′-triphosphate (H_2_MTP-P_3_). **(D)** Drug-repurposing workflow identifying common drugs of folB and folX, narrowing from 2,102 FDA-approved drugs to 20 candidates through *in silico* prediction.

### Analysis of proteins of the folate pathway to identify the targets

To systematically compare proteins in the folate biosynthesis pathway, we retrieved 16 pathway-associated protein sequences from UniProt and calculated all-against-all pairwise sequence identities, which were visualized as a heatmap (Figure 2A). This analysis revealed that folE, folX, and folB form the closest cluster within the pathway at the sequence level. Although a relatively high sequence identity was also observed between pabA and folM (∼50%), the folX–folB pair shared 24% sequence identity, consistent with their clustering within the grouped branch in the similarity matrix. These results highlight folX and folB as closely related proteins in the folate biosynthetic network based on primary sequence similarity.

**Figure 2.**
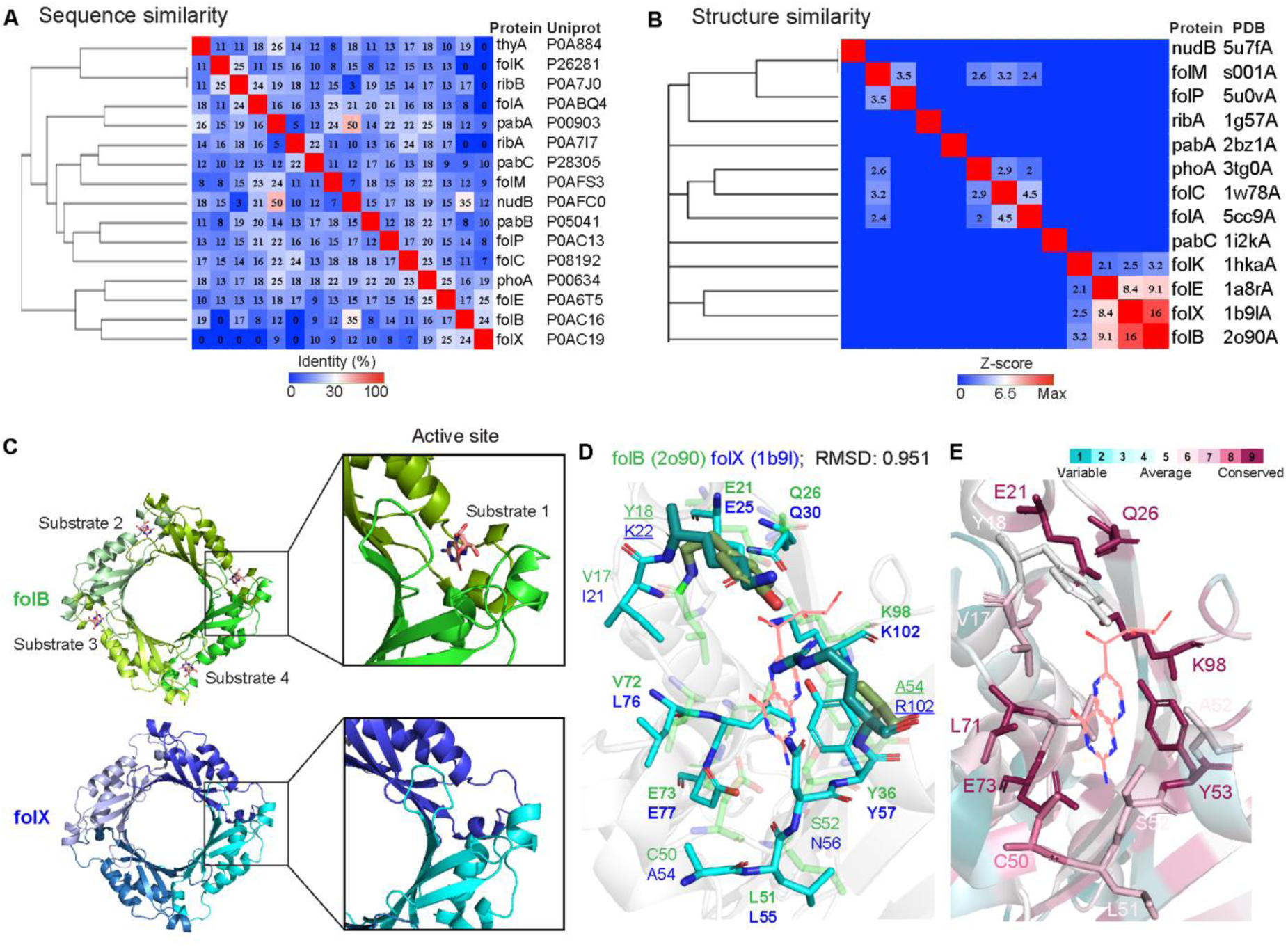
Multiple lines of evidence demonstrate that folB and folX are suitable co-targets within the folate biosynthesis pathway. **(A)** Hierarchical clustering of pairwise sequence identities for 16 pathway-associated proteins reveals a distinct folB–folX cluster. We calculated sequence identities using Clustal 2.1 via the UniProt portal. **(B)** Hierarchical clustering of pairwise structural similarity Z-scores, computed using DALI, shows that folB and folX are the most structurally similar proteins in the pathway. **(C)** Overall structural comparison of folB and folX tetramers, highlighting their highly similar quaternary arrangements and the identical location of the active sites at the dimer interfaces. **(D)** Superposition of the folB and folX binding pockets, yielding an RMSD of 0.951 Å. Residues in bold indicate identical amino acids, whereas residues underlined denote positions with different amino acid types. **(E)** Evolutionary conservation of the folB binding site, as analysed by ConSurf, demonstrating that most active-site residues display high conservation scores (grade >6).

Given that our drug screening strategy is structure-based, we next compiled all experimentally determined structures available in the PDB for proteins in the folate biosynthesis pathway and carried out a comprehensive pairwise structural comparison using the Dali server(Holm 2022). Consistent with our sequence-based analysis, folX and folB exhibited the highest degree of structural similarity among all proteins examined, with a Dali Z-score of 16 (Figure 2B), indicating a close structural similarity between these two enzymes. We further analysed folB and folX at the three-dimensional level, focusing on (i) the functional (substrate) site, (ii) the architecture of the binding pocket and its substrate binding residues, and (iii) residue conservation within the binding pocket. At the whole-protein level, both folB and folX adopt tetrameric assemblies, with the active sites located at the interface between two monomers (Ploom, Haußmann et al. 1999, Blaszczyk, Lu et al. 2014) (Figure 2C). Structural superposition of the ligand-binding pockets revealed a close match, with an RMSD of 0.951 Å, and the two enzymes largely sharing residues with equivalent physicochemical properties in their active sites—for example, folB: E21, Q26, K98, and Y53 correspond to folX: E25, Q30, K102, and Y57 (Figure 2D). Nevertheless, some notable differences in amino acid identity were observed in the binding pockets (underlined in Fig 2D), such as folB Y18 and A54 versus folX K22 and R102, respectively. The crystal structure of folB was solved in complex with neopterin (NEU), allowing us to directly examine the interaction between protein and ligand. We docked NEU into the folX structure (PDB: 1B9L) and observed highly similar binding interactions in both proteins (Supplementary Figure 1). In each case, the hydroxyl groups of NEU form multiple hydrogen bonds with active-site residues (folB: E21, V17, K98; folX: E25, R58, K102), while the amino (–NH₂) and carbonyl (C=O) groups on the pteridine ring also participate in hydrogen bonding with surrounding residues. In addition, NEU engages in van der Waals interactions with V17 in folB and Y57 in folX. Overall, these indicate a largely conserved ligand-binding pattern between folB and folX, while the differences in specific contacting residues point to subtle, and potentially functionally relevant, divergence between their active sites.

Finally, we assessed the evolutionary conservation of active-site residues using the ConSurf server(Yariv, Yariv et al. 2023). The analysis showed that most residues forming the folB active site are highly conserved, with ConSurf conservation grades > 6 (Figure 2E), indicating strong evolutionary pressure to maintain this pocket. Together with the high similarity between folB and folX, as well as their nearly identical binding environment, these findings support that folB and folX represent suitable co-targets for the rational design of dual inhibitors aimed at simultaneously modulating both enzymes in the folate biosynthesis pathway.

### Identification of dual inhibitors of folB and folX

To identify dual inhibitors of folB and folX, we carried out structure-based virtual screening using the molecular docking program GEMDOCK(Yang and Chen 2004), a docking tool based on a genetic algorithm, against a library of 2,102 FDA-approved drugs. Docking was performed with default binding-site settings, 70 evolutionary generations, a population size of 1,000, and a generation of 10 poses per compound (Figure 3A). For each target, the 3,000 best-scoring poses, ranked by GEMDOCK binding energy divided by the square root of N (N is the number of heavy atoms of the compound), were retained for further analysis. We then applied an additional post-docking scoring scheme, ComBX, to filter compounds that satisfy common interaction rules for both folB and folX. These rules require: (i) engagement of two predefined anchor regions (HV1 and HV2) (Figure 3B); (ii) overlap of key interacting residues with those involved in substrate recognition; (iii) consistent occurrence among the top 3,000 poses for both proteins; and (iv) favourable hydrogen-bond and van der Waals interactions with the residues listed in Figure 3C (see Methods for details).

**Figure 3.**
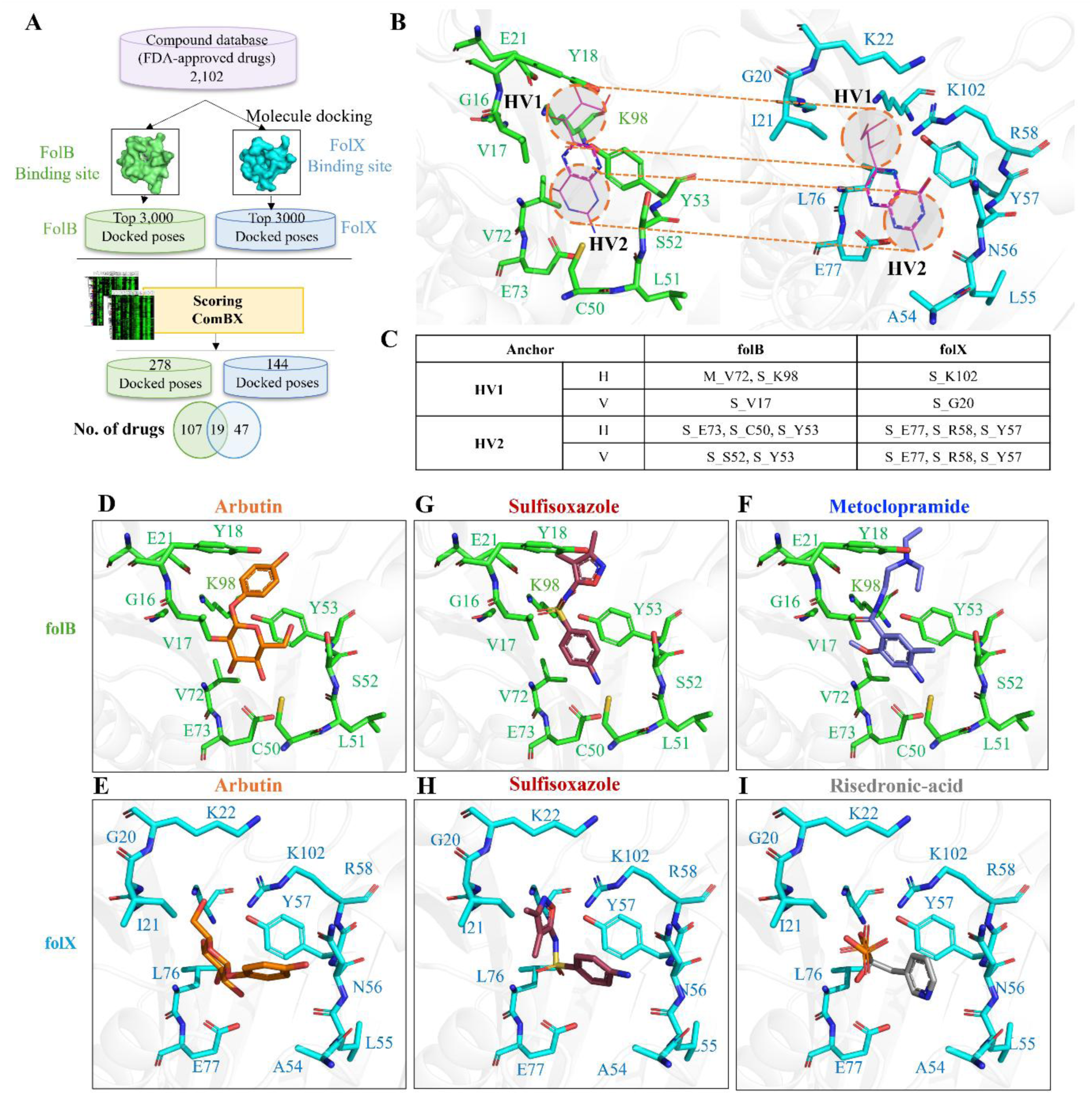
Structure-guided Repurposing of dual inhibitors targeting folB and folX. **(A)** Overview of the virtual screening strategy for folB and folX. The top 3,000 docked poses for each protein were selected and filtered based on common binding interactions. A total of 146 drugs were identified, including 19 common hits, 107 specifics to folB, and 47 specifics to folX. **(B)** Common binding interactions are shared by both folB and folX. HV1 and HV2 represent the location of the pocket required to satisfy both hydrogen-bond and van der Waals interactions. **(C)** List of anchor residues for both folB and folX. **(D** and **E)** Docked poses of representative compounds Arbutin on folB and folX, respectively. (**G** and **H**) Docked poses of representative compounds Sulfisoxazole on folB and folX, respectively. (**F**) Docked pose of Metoclopramide on folB. (**I**) Docked pose of Risedronic acid on folX.

Using the ComBX scoring method, we next analysed the filtered poses for each protein. Compounds showing in the top-ranking poses, suggesting stable accommodation within the active-site cavity, were retained as final candidates. This procedure yielded 278 docked poses corresponding to 126 distinct drugs for folB, and 144 docked poses corresponding to 66 distinct drugs for folX. Notably, 19 drugs were common to both sets, indicating that these molecules have the potential to bind both folB and folX, or preferentially bind only one of the two enzymes. Among the overlapping hits, Arbutin and Sulfisoxazole were selected as representative drugs predicted to target both folB and folX. The docking poses of Arbutin and Sulfisoxazole in folB are shown in Figure 3D and 3G, respectively, while their corresponding poses in folX are presented in Figure 3E and 3H. The β-D-glucose moiety of Arbutin engages in extensive van der Waals (vdW) and hydrogen-bonding interactions with residues in both active sites (folB: A70, V72, E73, A54, S52; folX: E77, L76, Y57, R58, E25). Its phenolic ring forms a π–π stacking interaction with Y53 in folB and strong van der Waals contacts with Y57 and K102 in folX (Supplementary Figure 2A and 2D). For Sulfisoxazole, the aniline ring participates in π–π interactions with Y53 in folB and Y57 in folX, further stabilizing its binding in the active sites of both enzymes (Supplementary Figure 2B and 2E). Metoclopramide was predicted to bind to folB only, with its π–π stacking interaction with Y53 in folB and a hydrogen bond with main chain V72 and E73 (Figure 3F and Supplementary Figure 2C). Risedronic acid was predicted to bind to folB with its phosphate group forming strong hydrogen bond interactions to residues R58, K102, Y57, and E77 in folX (Figure 3I and Supplementary Figure 2F).

Next, we utilized compound similarity analysis to identify representative drugs from each cluster. Ultimately, we selected 20 drugs for further *in vitro* evaluation: eight from the distinct set of folB, seven from the intersection set, and five from the distinct set of folX.

### Arbutin exhibits a folate-linked inhibitory phenotype

We first quantified growth inhibition across six mechanistically diverse putative antimicrobials. The compounds inhibited bacterial growth with IC₅₀ values ranging from 0.08 to 0.27 μg/ml, indicating potent antibacterial activity across all hits (Figure 4A). Exogenous folate supplementation produced a pronounced and concentration-independent rescue of arbutin-mediated growth inhibition (Figure 4B), indicating that arbutin perturbs a metabolic pathway that is either directly dependent on, or functionally coupled to, folate availability. Folate supplementation experiments were carried out for cells under Gemcitabine, Valacyclovir and Cefanoxime treatments (Supplementary Figure 3). Absence of folate rescue under these treatment conditions indicates that these compounds do not perturb folate biosynthesis, and therefore their molecular targets likely lie outside the folate pathway. Sulfisoxazole exhibited the expected partial rescue (Figure 4C), reflecting its PABA-competitive inhibition in early folate biosynthesis. Cefanoxime, a non-folate-associated β-lactam, showed no folate dependence (Supplementary Figure 3). These signatures collectively delineate arbutin as a compound whose growth-inhibitory effect is mechanistically proximal to folate metabolic flux rather than global nucleotide depletion or cell-wall biosynthesis defects.

**Figure 4.**
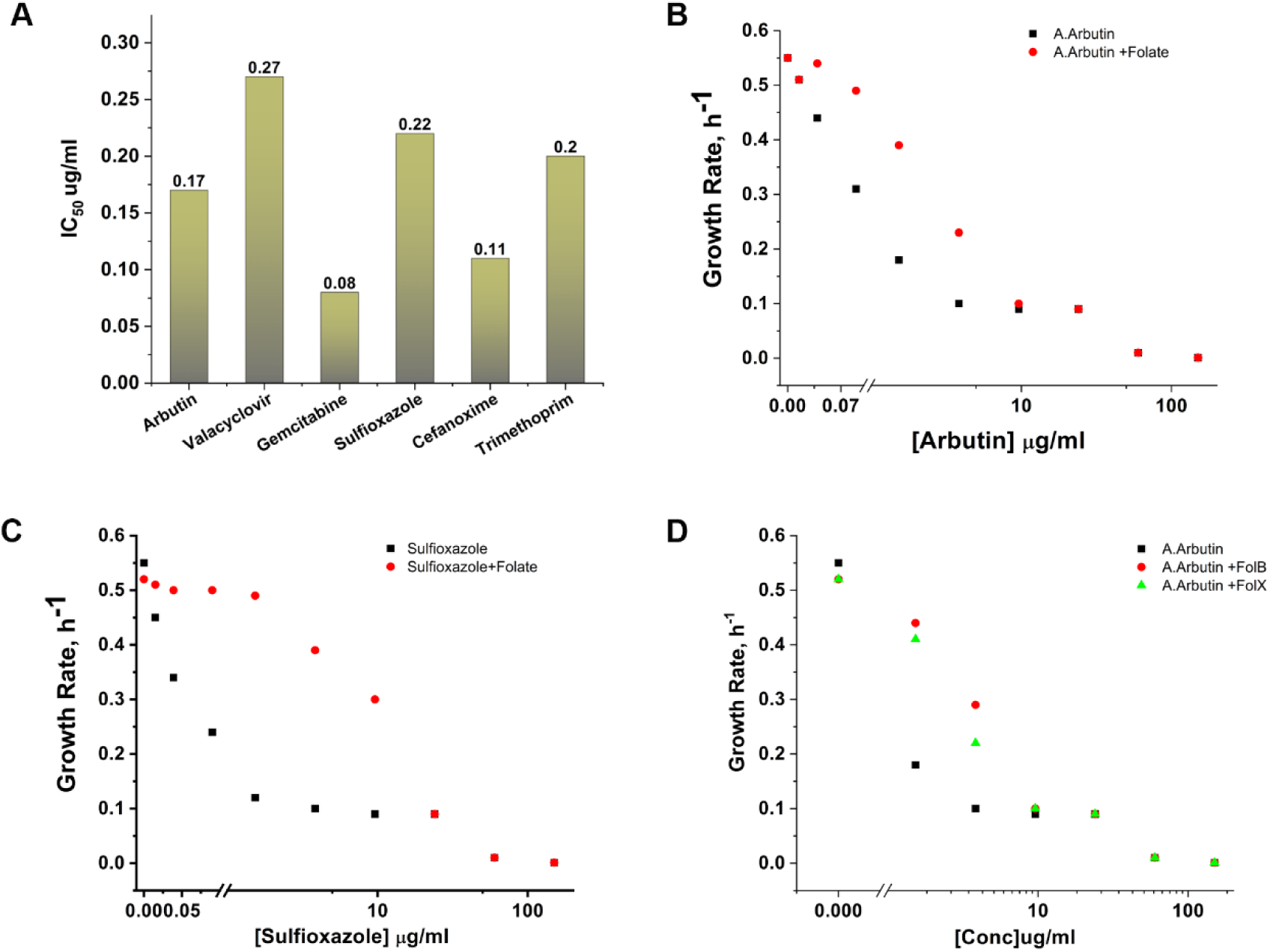
Folate supplementation experiments show arbutin co-targets folB and folX. (A) Cell Growth Inhibition Profile. IC₅₀ values for arbutin, valacyclovir, gemcitabine, sulfisoxazole, cefanoxime, and trimethoprim measured under standard growth conditions. Bars represent mean IC₅₀ (µg/mL), highlighting relative potencies across compounds. **(B)** Effect of folate supplementation on Arbutin-mediated growth inhibition. Bacterial growth rates are plotted as a function of increasing Arbutin concentration in the absence (black) and presence (red) of exogenous folate. Partial restoration of growth upon folate addition indicates that Arbutin disrupts folate biosynthesis. **(C)** Folate-dependent rescue of growth inhibition by Sulfisoxazole. Growth rates measured across increasing Sulfisoxazole concentrations show partial recovery with folate supplementation, consistent with inhibition of folate metabolism. **(D)** Gene-specific complementation analysis under Arbutin treatment. Growth rates are shown for cells expressing folB (red) or folX (green) compared to control (black) across increasing drug concentrations.

We next examined whether folB and folX contribute to the cellular response to arbutin. To test this, we performed gene-dosage rescue experiments using strains overexpressing either gene. folB encodes dihydroneopterin aldolase, while folX encodes dihydroneopterin triphosphate epimerase. Expression of either gene restored bacterial growth in the presence of arbutin (Figure 4D), with folB providing a stronger rescue effect, particularly at intermediate arbutin concentrations. These findings demonstrate that increasing the cellular abundance of folB or folX mitigates arbutin-mediated growth inhibition, implicating both enzymes in the arbutin-sensitive phenotype. Taken together with the docking analysis and folate supplementation experiments, these overexpression results support a model in which arbutin-induced growth inhibition arises from inhibitory interference with folB and folX function within the folate biosynthesis pathway.

To test whether genetic rescue reflects direct inhibition of folX and folB by arbutin, we quantified arbutin–protein interactions using amide-I-resolved Raman spectroscopic titrations. To enable comparison across different proteins and experimental conditions, we normalized Raman spectral intensities and scaled (log-transformed where appropriate) to account for differences in absolute signal magnitude. This scaling preserves relative binding trends while allowing consistent visualization across datasets. The interaction-specific amide I peak area increased progressively with ligand concentration (Supplementary Figure 4A, B) and approached saturation at higher concentrations, consistent with a concentration-dependent binding process. Hyperbolic fitting of the spectral response yielded apparent dissociation constants (K_d_). Because Raman spectroscopy captures ligand-induced structural and environmental perturbations rather than direct binding thermodynamics, the derived K_d_ values represent effective interaction parameters reporting on qualitative effect of binding rather than equilibrium binding affinities. As a reference, TMP displayed a clear saturable interaction profile, with nonlinear fitting yielding an apparent dissociation constant (Kd) of 9.01 μM (Supplementary Figure 5). Similarly, arbutin showed comparable saturable binding behaviour with folB and folX, with fitted Kd values of 13.68 μM and 10.55 μM respectively (Supplementary Figure 4A, B). Although these values may differ from affinities obtained using thermodynamic techniques such as ITC due to the indirect nature of the affinity readout in Raman spectroscopy, the comparable apparent low-micromolar range observed for TMP and Arbutin indicates that the Raman-derived signal shows specific ligand engagement relevant to functional inhibition. Notably, these effective Kd values are consistent with the in vivo IC₅₀ measurements (arbutin: 0.08-0.27 μg/mL ≈ 0.3-1.0 μM), supporting the physiological relevance of the observed binding interactions.

Our growth experiments indicate Arbutin’s inhibitory phenotype is strongly folate responsive.

### Dual-target Arbutin imposes strong evolutionary constraints, unlike folB or folX specific inhibitors

We monitored how *E. coli* populations adapted to increasing concentrations of Arbutin, Metoclopramide, and Risedronic acid during 41 days of serial passaging (Figure 5A). We selected Metoclopramide and Risedronic acid as inhibitors specific to folB and folX, respectively, in order to directly compare adaptation under single-node versus dual-node inhibition within the folate pathway. The trajectories diverged rapidly. Under Metoclopramide and Risedronic acid, the populations tolerated progressively higher concentrations almost throughout the entire experiment, reaching ∼ 4-5 times the WT IC₅₀ by day 41. In contrast, the IC_50_ along Arbutin trajectories increased only gradually and plateaued near 1.5× IC₅₀. These trends show that single-target inhibition at folB or folX does not prevent continuous adaptive gains, whereas dual folB/folX inhibition sharply restricts the evolutionary increase in tolerated concentration.

**Figure 5.**
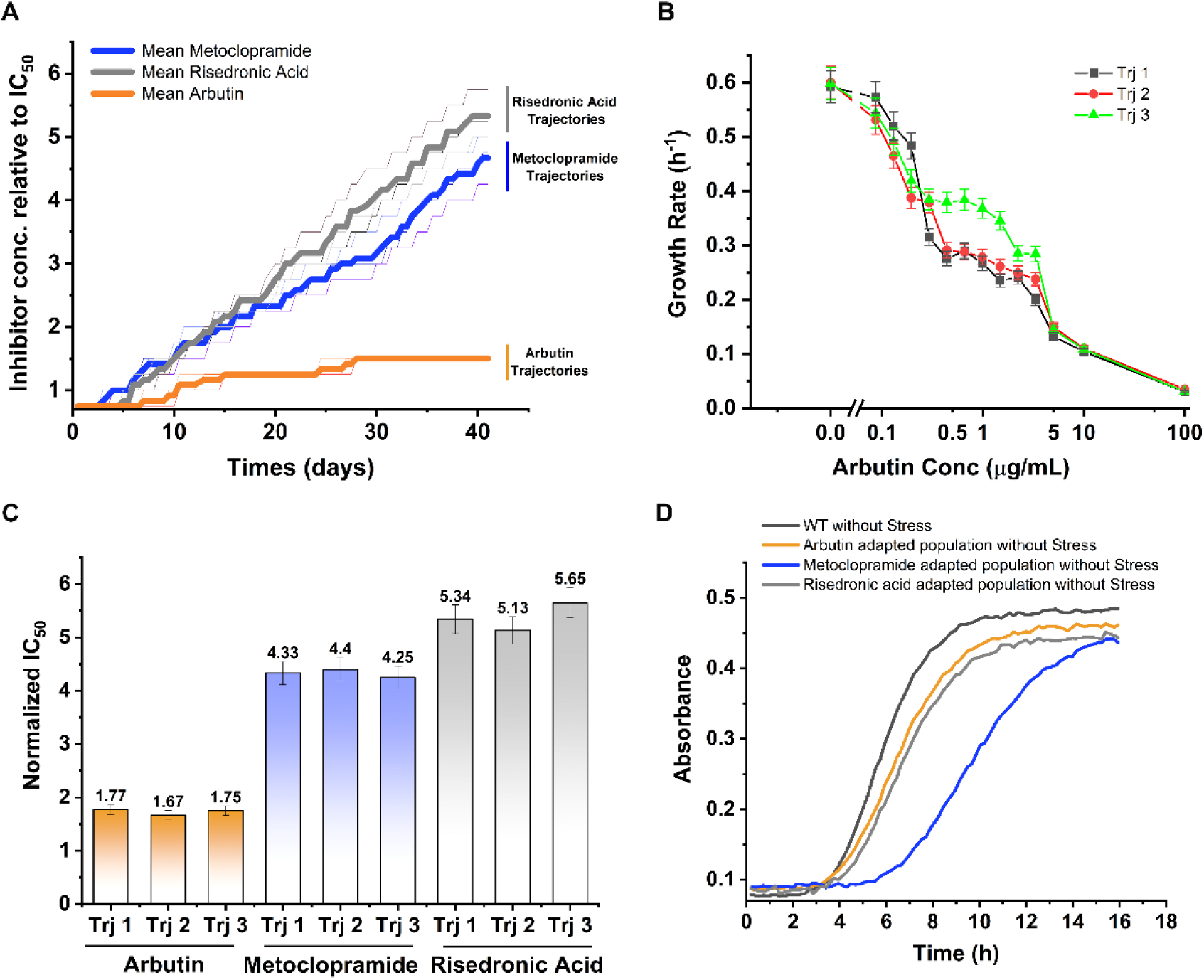
Evolutionary responses of E. coli to dual and single-target inhibition. (A) Changes in tolerated drug levels over 41 days of evolution. E. coli populations evolved in the presence of Arbutin (blue; dual folB/folX inhibitor), Metoclopramide (grey; folB inhibitor), or Risedronic acid (red; folX inhibitor). We plotted drug concentrations relative to the WT IC₅₀. Populations evolving in Metoclopramide and Risedronic acid steadily increased their tolerated concentrations, while Arbutin-evolved populations showed only gradual increases and plateaued near ∼1.5 times IC₅₀. Curves represent mean values from three independent trajectories. (B) Growth-rate measurements for Arbutin-evolved populations across an Arbutin concentration gradient. Growth rates declined with increasing drug concentration in all trajectories. Symbols represent three independent evolution lines; error bars denote standard deviation from triplicate growth assays. (C) Normalized IC₅₀ values for evolved populations. Arbutin-evolved trajectories showed modest increases relative to the ancestor, whereas Metoclopramide- and Risedronic acid–evolved trajectories exhibited large increases in IC₅₀. Bars represent mean values for each trajectory. (D) Drug-free growth curves for the ancestral strain and evolved populations. All evolved populations showed slower growth and lower final densities relative to the ancestral strain. Curves represent mean absorbance measurements from biological replicates.

To quantify physiological tolerance to Arbutin, we measured growth rates across a concentration gradient for all three evolved trajectories(Zhang, Chowdhury et al. 2021) (Figure 5B). The growth rate of each evolved population decreased steeply as Arbutin concentration increased. While the evolved populations exhibited modest gains in growth rate at intermediate Arbutin concentrations, these changes did not substantially alter the inhibition pattern, indicating that the adaptations provided only minor physiological relief. This pattern demonstrates that, despite prolonged selection, the Arbutin-evolved populations retain limited ability to grow at elevated drug levels. In contrast, growth-rate measurements for the folB-evolved trajectories (Supplementary Figure 6A) and folX-evolved trajectories (Supplementary Figure 6B) show less pronounced decreases in growth rate with increasing drug concentration. Both single-target conditions supported substantially higher growth at intermediate and high concentrations, highlighting their ability to maintain physiological performance at drug levels that strongly inhibit the Arbutin-adapted populations.

We next calculated IC₅₀ values for all evolved populations (Figure 5C). For Arbutin, evolved trajectories showed only a small increase in IC₅₀, ranging from 1.6 to 1.8 times the WT value. In contrast, trajectories evolved under Metoclopramide and Risedronic acid showed substantially higher IC₅₀ values, approximately from 4 to 4.5 times for folB-only inhibition and 5.1 to 5.6 times for folX-only inhibition. These differences are consistent with the adaptive trajectories in Fig. 5A, confirming that dual-target inhibition constrains resistance, while single-target inhibition permits pronounced shifts in drug sensitivity. IC₅₀ measurements further emphasize that folX-only-evolved populations reached IC₅₀ values of 5.9-6.4 µg/mL (Supplementary Figure 6C), and folB-only–evolved lines reached 5.65-5.9 µg/mL (Supplementary Figure 6D), both far exceeding the modest increases observed under Arbutin treatment.

Finally, we compared the growth curves of the ancestral strain with those of evolved populations in the absence of drug (Figure 5D). The ancestral strain grew rapidly and reached a high final cell density. All evolved lines displayed slower growth, with reduced maximal absorbance, indicating that long-term adaptation carries a measurable fitness cost. The Arbutin-adapted populations grew more slowly than wild type, and the Metoclopramide- and Risedronic acid–adapted lines showed even more pronounced reductions, suggesting that resistance acquisition is associated with trade-offs in basal growth.

Together, these results show that *E. coli* responds very differently to inhibition at distinct nodes of the folate pathway. Dual inhibition of folB and folX through Arbutin leads to slow adaptation, minimal increases in IC₅₀, and highly similar phenotypic trajectories across three independent populations, characterized by early plateauing of IC₅₀ values and limited adaptive gains over time. In contrast, single-target inhibition at folB or folX allows rapid and repeated gains in resistance, reflected in both elevated growth rates under stress and substantially higher IC₅₀ values. These observations indicate that simultaneous targeting of multiple folate enzymes sharply limits the accessible mutational routes for resistance, while single-enzyme inhibition leaves broader adaptive paths available.

### Genome-wide analysis reveals condition-dependent gene duplication under folB/folX inhibition

We performed whole-genome sequencing on three independently evolved *E. coli* populations exposed to Arbutin, the dual folB/folX inhibitor (mentioned as Trj1, Trj2, Trj3 in Figure 6A), using the native BW27783 strain as the reference(Bhattacharyya, Bershtein et al. 2016). We observed both large-scale chromosomal duplications and point mutations in populations evolved under Arbutin and folX-specific inhibition by Risedronic acid, whereas FolB-specific inhibition by Metoclopramide resulted exclusively in point mutations without any detectable duplication (Figure 6A).

**Figure 6.**
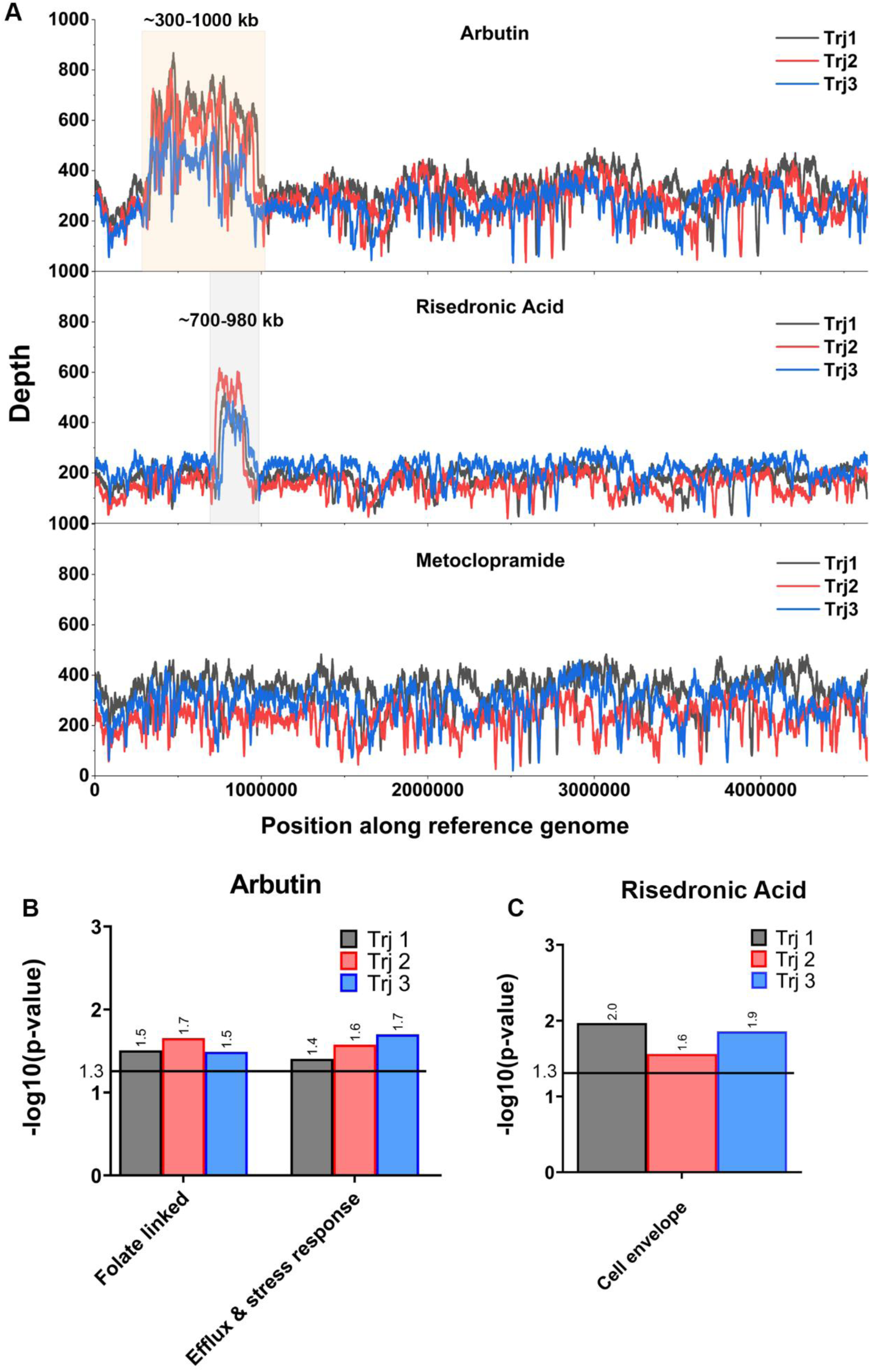
Genome-wide duplication patterns and functional composition of amplified regions under drug selection. (A) Genome-wide sequencing depth profiles of evolved *E. coli* populations selected under Arbutin, Risedronic acid, or Metoclopramide. Sequencing depth (y-axis) is plotted as a function of genomic position along the reference genome (x-axis). Traces correspond to three independent evolutionary trajectories (Trj1-Trj3). Shaded regions indicate reproducibly amplified genomic segments characterized by elevated read depth relative to the genomic background, consistent with large-scale segmental duplications. (B and C) Functional enrichment analysis of genes within duplicated genomic regions identified in evolved populations. Enrichment significance is reported as −log₁₀(p-value), calculated by comparing the frequency of genes within each functional category in the duplicated region relative to the genomic background. The horizontal line indicates the significance threshold (p=0.05; −log₁₀(p) ≈ 1.3). For Arbutin-evolved populations (B), duplicated regions show consistent enrichment of folate-linked and efflux and stress-response gene categories across independent trajectories, indicating amplification of functions associated with one-carbon metabolism and cellular stress buffering. For Risedronic acid-evolved populations (C), enrichment is observed primarily in cell envelope-associated genes, suggesting a distinct functional bias in the duplicated segment. Bars represent independent evolutionary trajectories (Trj1, Trj2 and Trj3). Differences in bar height reflect relative enrichment significance and should be interpreted qualitatively rather than as direct statistical comparisons between categories or trajectories.

All Arbutin-evolved trajectories exhibited a prominent large-scale chromosomal gene duplication spanning approximately from 300 to 1000 kb (Figure 6A). The sequencing depth profiles were highly consistent across replicates and showed a stable ∼3 to 4-fold increase in copy number across this region, indicating a reproducible genome-dosage change. In addition to this duplication, each trajectory accumulated point mutations distributed across the genome, demonstrating that adaptation proceeds through a combination of local mutational changes and large-scale genomic amplification under dual inhibition.

To determine whether this duplication reflects a general response to antifolate stress or a condition-specific outcome, we analysed populations evolved under Metoclopramide (folB-specific) and Risedronic acid (folX-specific) (Figure 6A). Risedronic acid-evolved populations exhibited both point mutations and a narrower, lower-amplitude chromosomal duplication localized in the ∼700-980 kb region. Whereas, under Metoclopramide, sequencing-depth profiles remained flat across all trajectories, with no detectable increase in coverage in any genomic region. All observed genetic changes consisted of point mutations, small insertions or deletions, or localized rearrangements, indicating that adaptation proceeds through localized sequence variation without engaging genome-scale amplification. The coexistence of mutations and duplication under both Arbutin and Risedronic acid folX inhibition indicates that genome-dosage changes are selectively engaged under these conditions, whereas folB-only inhibition is accommodated through localized mutational adjustments without detectable large-scale genomic restructuring. The emergence of large chromosomal duplications is consistent with a higher fitness cost, as increased gene dosage imposes additional replication and transcriptional demands on the cell.

To further characterize the functional content of the duplicated regions, we performed enrichment analysis of gene categories within amplified segments relative to the genomic background. In order to determine whether the observed enrichment is specific to adaptation or reflects a general gene composition bias within the *E. coli* genome, we quantified enrichment of specific genes in a duplicated region vs null model representing compositional distribution of genes in random samples of genomic regions of the same length. To that end we compared the observed fraction of genes in each functional category within duplicated regions to a null distribution generated from 10,000 randomly sampled genomic segments of *E. coli* chromosome of identical length (see Methods). This analysis quantifies whether specific functional classes are overrepresented within duplicated regions and thereby identifies the cellular processes preferentially expanded during adaptation.

In Arbutin-evolved populations, the duplicated region showed consistent highly statistically significant enrichment of folate-linked genes and efflux and stress-response functions across all trajectories (Figure 6B). Here, folate-related genes include both canonical folate biosynthesis genes and additional genes linked to one-carbon and nucleotide metabolism (see Methods). Folate-related genes were significantly enriched in the duplicated region across Arbutin-evolved trajectories (Trj1: p-value = 0.032, Trj2: p-value = 0.02, Trj3: p-value = 0.032), along with efflux and stress-response genes (Trj1: p-value = 0.04, Trj2: p-value = 0.03, Trj3: p-value = 0.02). These enrichments indicate non-random expansion of pathway-relevant functions under dual inhibition. Notably, enrichment of folate-related genes is directly relevant to the mechanism of inhibition, as Arbutin simultaneously targets folB and folX within the folate biosynthesis pathway. Increased gene dosage of downstream folate-pathway components can help sustain one-carbon flux and partially compensate for reduced enzymatic activity at these inhibited steps. In addition, enrichment of efflux and stress-response genes suggests an expanded capacity to mitigate intracellular drug accumulation and buffer redox and proteostatic stress associated with pathway disruption.

In Risedronic acid-evolved populations, cell envelope associated genes were primarily enriched in the duplicated region (Trj1: p-value = 0.01, Trj2: p-value = 0.025, Trj3: p-value = 0.013). (Figure 6C). This enrichment was also statistically significant relative to random expectation (p < 0.05), supporting a condition-specific adaptive response. Amplification of these genes may alter membrane composition or permeability and potentially reduce intracellular drug access or modifying stress signalling at the cell boundary. Together, these patterns indicate that distinct functional classes are preferentially amplified under different inhibition conditions, reflecting differences in how cells accommodate pathway perturbation.

### Target-specific fitness constraints define mutational trajectories and adaptive solutions in evolving populations

Whole-genome sequencing of Arbutin, Risedronic Acid, and Metoclopramide evolved populations revealed distinct adaptive strategies that reflect differences in evolutionary constraint at the functional level. We classified all mutated genes into gene families to compare trajectories at the functional level (Supplementary Table 2 to 13). The number and recurrence of gene families varied across conditions, indicating differences in accessible evolutionary paths and associated fitness costs.

Under Arbutin selection, populations gained fitness through multiple genetic routes with limited gene-level overlap between trajectories, except for pphB in Trj1 and Trj3 (Supplementary Table 5). However, trajectories repeatedly targeted similar gene families, including DNA repair, rRNA methylation, and operon-level systems such as uid and ygf.

Within trajectories, mutations clustered in metabolic families such as sda and tdc, indicating pathway-level rewiring. These patterns indicate that Arbutin permits multiple adaptive routes while constraining outcomes at the level of functional processes, consistent with a moderate fitness cost and partial restriction of evolutionary trajectories.

Different resistance trajectories evolving under folX-only inhibition, showed no shared genes between Trj1-Trj3 (Supplementary Table 5 to Supplementary Table 8). Instead, populations improved fitness through distinct mutations that converged only at the gene family level. Repeated targeting of IS elements, yed-family genes, and pathway-specific families such as mgl, hyb, and frl indicates distributed adaptation (Supplementary Table 9). This pattern indicates that selection acts across a broad functional landscape, enabling multiple independent adaptive routes with minimal constraint at the gene level, consistent with a high but diffuse fitness cost.

In contrast, evolution of resistance under folB-only inhibition drove strong and repeatable adaptation. Trj2 and Trj3 repeatedly acquired mutations in the same set of genes, including glcD, cysI, hisB, yejM, ydjY, gspL, insI1, mntR, ddpX, yaiO, and wcaI (Supplementary Table 10 to Supplementary Table 13). These genes belong to a limited set of families involved in metabolism, envelope structure, secretion, and regulation. This tight recurrence indicates that adaptation is restricted to a narrow set of accessible solutions, reflecting strong constraint on evolutionary trajectories and a focused fitness cost landscape.

Together, these results define a continuum of evolutionary constraints across conditions that reflects differences in the underlying fitness landscape. Arbutin allows flexible, partially convergent adaptation. folX drives highly distributed responses with only functional-level convergence. folB restricts adaptation to specific genes and pathways. This progression demonstrates that the nature of metabolic perturbation determines how fitness costs shape the accessibility and diversity of evolutionary trajectories, ranging from flexible network-level adjustments to highly constrained, repeatable solutions.

## DISCUSSION

A large fraction of antibacterial agents targets metabolic enzymes, reflecting the central role of metabolism in sustaining cellular growth and replication(Stokes, Lopatkin et al. 2019). Enzymes within core biosynthetic pathways are particularly attractive targets because they control flux distribution through essential processes such as nucleotide production, amino acid synthesis, and redox balance(Walsh 2003, Brown and Wright 2016). Inhibiting such enzymes imposes immediate and system-wide constraints on cellular physiology, as metabolic networks operate under tightly coupled flux dependencies(Zampieri, Enke et al. 2017). However, despite their essentiality, metabolic targets often remain vulnerable to rapid resistance evolution, as mutations or regulatory changes can impact metabolic flux, compensate for pathway perturbations, or alter drug-target interactions without compromising enzyme function(Palmer and Kishony 2013, Zampieri, Enke et al. 2017).

The folate biosynthesis pathway is widely targeted by antibiotics but remains prone to rapid resistance evolution(Bertacine Dias, Santos et al. 2018). This pathway supplies tetrahydrofolate cofactors required for one-carbon transfer reactions that govern nucleotide biosynthesis, amino acid metabolism, and methylation reactions across bacteria and many lower eukaryotes(Ducker and Rabinowitz 2017). Inhibiting folate metabolism directly constrains DNA replication and cell proliferation, yet resistance arises rapidly through binding-site mutations, gene amplification, or metabolic rewiring that restores pathway flux(Huovinen, Sundström et al. 1995, Toprak, Veres et al. 2012, Rodrigues, Bershtein et al. 2016, Rodrigues and Shakhnovich 2019). These observations highlight a key limitation: Inhibition of a single metabolic node perturbs flux but still allows for compensatory adaptation. Here, we show that targeting folB and folX together restricts both the extent of resistance and the set of adaptive trajectories available to the population.

We demonstrate that coordinated inhibition of structurally convergent enzymes imposes a strong evolutionary constraint on bacterial adaptation. By integrating computational docking, biochemical validation, and long-term evolution experiments, we show that simultaneous perturbation of two structurally and functionally linked enzymes within a metabolic pathway alters the adaptive landscape, limiting the accessibility of mutational escape routes(Weinreich, Delaney et al. 2006). As a result, adaptation is redirected toward higher-cost, system-level responses rather than local target-specific changes.

Consistent with this framework, our study shows that resistance depends not only on inhibitory potency but on how inhibition reshapes the accessibility of adaptive states within the fitness landscape. Under single-target inhibition, resistance emerges readily through multiple low-cost mutational paths that preserve function while reducing drug binding. In contrast, simultaneous perturbation of two functionally linked enzymes introduces correlated constraints that eliminate these low-cost routes(Kryazhimskiy, Rice et al. 2014, Szili, Draskovits et al. 2019). As a result, populations are forced toward rarer, higher-cost adaptive solutions, leading to slower and incomplete resistance acquisition(Lukačišinová, Fernando et al. 2020).

Our experimental evolution data directly support this model by revealing distinct adaptive outcomes under different inhibition conditions. Under single-enzyme inhibition (Metoclopramide and Risedronic acid inhibition), populations rapidly increased tolerance (∼ 4-5-fold) with consistent trajectories across replicates. In contrast, under dual inhibition, populations showed only a modest increase in tolerance (∼1.5-2-fold) and remained sensitive across the drug gradient, indicating a constrained adaptive response. Whole-genome sequencing reveals clear differences in adaptive responses across conditions. Under folB inhibition (Metoclopramide), populations acquire a restricted set of recurrent mutations. Under folX inhibition (Risedronic acid), populations accumulate diverse mutations with convergence at the functional level, often accompanied by chromosomal duplication. In contrast, under dual inhibition (Arbutin), populations follow multiple genetic routes but remain constrained at the pathway level, with adaptation involving both mutations and genome-scale duplication. Together, these results indicate that resistance outcomes are governed by the distribution and fitness cost of accessibility and cost of adaptive solutions rather than inhibitory strength alone(Toprak, Veres et al. 2012).

Mechanistically, these differences reflect how populations balance localized and genome-level responses under selection. Point mutations provide targeted, low-cost adjustments, whereas chromosomal duplication enables rapid increases in gene dosage across multiple loci, offering a broader compensatory response. Under folX and dual inhibition, selective pressure is not fully resolved by localized changes alone, leading to the engagement of both mutational and genome-level responses. In contrast, folB inhibition is effectively accommodated through gene-specific mutations, reducing the need for large-scale genomic changes. This suggests that duplication is selected when low-cost mutational solutions are insufficient, while mutations refine adaptation once viable solutions are established.

These findings indicate that when structurally convergent enzymes are co-targeted, the system cannot compensate through independent, target-specific changes. This limits the range of viable adaptive responses and shifts adaptation toward system-level reorganization.

The present study focuses on a single metabolic pathway in one organism, allowing us to directly investigate how coordinated perturbation modulate adaptive responses. In this framework, genome-scale duplications and mutational responses reveal how adaptation proceeds under selection, while the long-term stability, associated fitness costs, and reversibility of these adaptive states remain to be determined. More broadly, this work reframes antimicrobial design as the problem of shaping evolutionary outcomes, where the objective is not only to inhibit growth but to restrict the range of viable adaptive responses available to the organism. Simultaneous targeting of functionally linked enzymes therefore provides a rational strategy to limit resistance evolution by restricting accessible evolutionary routes.

This work focuses on a single organism and a defined metabolic pathway. The generality of these findings across other bacterial systems, including clinically relevant ESKAPE pathogens(Santajit and Indrawattana 2016, De Oliveira, Forde et al. 2020), remains to be established. The analysis is restricted to the folate biosynthesis pathway, and whether similar constraints arise across other metabolic networks remains to be tested. Incorporating transcriptomic, proteomic, and metabolite-level measurements will capture additional layers of adaptive response, including dynamic reorganization of the resistome(Lopatkin, Stokes et al. 2019). Cellular drug uptake, intracellular partitioning, and effects on secondary metabolism may further influence fitness under selection and remain to be investigated(Richter, Drown et al. 2017, Cama, Voliotis et al. 2020). The observed adaptive responses may also depend on compound-specific properties and require validation across different inhibitor classes and pathways(Palmer and Kishony 2014).

Future work should integrate metabolic modelling with isotope-based tracking to quantify changes in flux across metabolic pathways to quantify system-level metabolic responses under coordinated inhibition. Combining multi-omics approaches with these analyses and evolutionary experiments will help define the co-targeted resistome at higher resolution. Optimizing dual-target scaffolds such as Arbutin and testing them across additional pathway contexts may further strengthen evolutionary constraint. Combining such strategies with efflux inhibitors or existing antibiotics may also reduce residual adaptation.

## Materials and Methods

### Screening Of the Folate Synthesis Pathway

Proteins annotated in the folate biosynthesis pathway (KEGG map00790) were collected from the KEGG Pathway Database. Protein sequences were retrieved from the UniProt database, and pairwise sequence identities were calculated using CLUSTAL 2.1 implemented within UniProt (https://www.uniprot.org/). Protein structures with experimentally solved coordinates were retrieved from the Protein Data Bank (PDB)(Burley, Berman et al. 2022) for further structural analysis. Individual PDB chains were extracted, and pairwise structural alignments were performed using Dali(Holm and Park 2000). The resulting pairwise Z-scores were used to construct a clustering tree with the Morpheus web-based clustering tool (https://software.broadinstitute.org/morpheus/), applying average linkage.

### Preparation Of Protein Structures, Compound Datasets and Docking

For both folB and folX, the representative structures were downloaded from the PDB, selected and aligned, where floX (PDB id: 1b9l) and floB (PDB id: 2o90)(Ploom, Haußmann et al. 1999, Blaszczyk, Lu et al. 2014). The 10 Å residues were selected around the bound ligand and were extracted to form the active site cavities for each protein. The binding sites were extracted using the SwissPDB viewer. The binding site cavities were then prepared for docking by defining the residue atom types and assigning charges using the GEMDOCK method. For the 2,102 FDA drug set, the FDA drug list assigned as “approved” by the Drugbank (https://www.drugbank.ca/) was collected, referencing our previous paper(Pathak, Chen et al. 2020), whose 3D structures were downloaded from ZINC15 (https://zinc15.docking.org/). The parameter of GEMDOCK was set with the settings of 70 generations, a population of 1,000, and 10 poses for each compound.

### Generation of Interaction Profiles

We then generate interaction profiles from molecular complexes and molecular interface families. In total, two interaction profiles are generated, including protein-ligand and protein-compound interactions. An interaction profile P is presented as

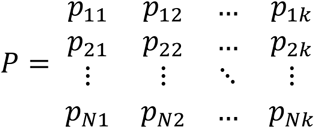

where *p*_*ij*_ is a binary value for molecular interactions. *p*_*ij*_ is set to 1 if the part of the molecule forms electrostatic, hydrogen-bonding, or van der Waals interactions with the protein subpockets; otherwise, *p*_*ij*_ is set to 0.

### Virtual Screening and Scoring for Identifying Dual Inhibitors

The current virtual drug screening used a docking-based approach, involving the filtering of drug poses using a common folB and folX (ComBX) scoring, *S*_ComBX_(*p*), where *p* is the top poses and defined in equation (1). We applied the docking tool GEMDOCK. In the virtual screening for drug repurposing, the set of 2,102 FDA-approved drugs was docked into each cavity using the specified iGEMDOCK parameters. The top 3,000 poses by energy are divided by the number of heavy atoms of the compound. For each protein were then selected for further analysis. The top 3,000 docking poses for each target were first ranked by the ComBX score and then filtered using predefined anchor-based interaction criteria. Core interactions were evaluated for two anchor regions, HV1 and HV2, and a pose was considered to satisfy an anchor only when all required interaction types and energy thresholds were met. In the interaction notation, S and M denote side-chain and main-chain contacts, respectively.

For FolB, a pose was considered to satisfy the HV1 anchor if it formed at least one hydrogen-bond (HB) interaction with either S_K98 or M_V72, with an interaction energy of ≤ −3 kcal/mol, and formed a Van der Waals bond (VB) interaction with S_V17 of ≤ −4 kcal/mol. A pose was considered to satisfy the HV2 anchor if it formed hydrogen-bond interactions with at least two of the following residues: S_E73 (≤ −4 kcal/mol), S_C50 (≤ −2 kcal/mol), and S_Y53 (≤ −4 kcal/mol), and also formed V-bond interactions with S_Y53 (≤ −15 kcal/mol) and M_S52 (≤ −1.5 kcal/mol).

The criteria were tailored into FolB anchor score for each pose *p,*

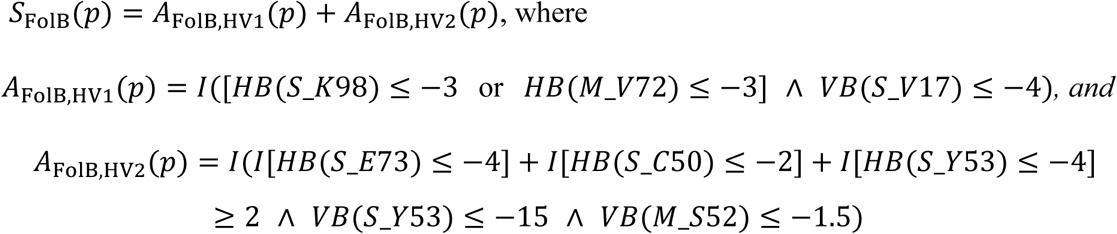

For FolX, a pose was considered to satisfy the HV1 anchor if it formed a hydrogen-bond interaction with S_K102 of ≤ −3.5 kcal/mol and a V-bond interaction with M_G20 of ≤ −1.5 kcal/mol. A pose was considered to satisfy the HV2 anchor if it formed hydrogen-bond interactions with at least two of the following residues: S_E77 (≤ −4 kcal/mol), S_Y57 (≤ −3 kcal/mol), and S_R58 (≤ −2.5 kcal/mol), and also formed V-bond interactions with S_E77 (≤ −3 kcal/mol), S_Y57 (≤ −6 kcal/mol), and S_R58 (≤ −2.5 kcal/mol).

The criteria were tailored into FolX anchor score for each pose *p*,

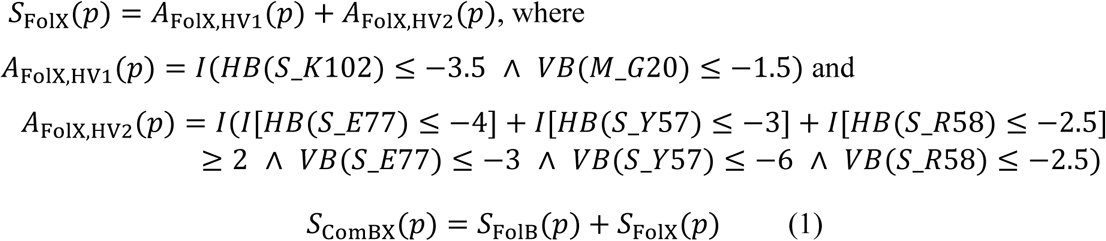

where *A*_FolB,HV1_, *A*_FolB,HV2_, *A*_FolX,HV1_, *A*_FolX,HV2_ ∈ {0,1}, and therefore *S*_FolB_(*p*), *S*_FolX_(*p*) ∈ {0,1,2} and *S*_ComBX_(*p*) ∈ {0,1,2,3,4}.

The pose was then classified as:

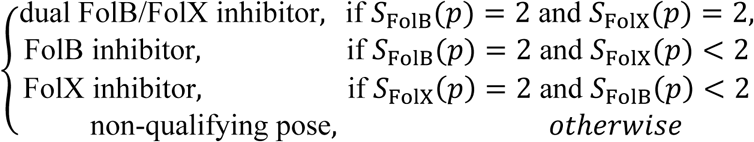

If the pose predicted as inhibitor, indicative of stabilized binding in the active site cavity, were selected for the final list of drug candidates. A total of 278 docked poses across 126 different drugs were filtered from folB, and 144 docked poses across 66 different drugs were filtered from folX. There is an overlap of 19 drugs between these two proteins. The overlapping compounds were predicted to act as dual inhibitors of both folB and folX, whereas the remaining compounds were identified as specific inhibitors for either folB or folX individually.

### Compound Features and Clustering

The complexed ligand structures were analysed and counted for the ligand 2D fingerprint features, a combination of structural features from Checkmol (with 204 predefined functional groups) and atomic composition (with 10 atom types to describe a ligand: carbon in ring, other carbon, nitrogen in ring, other nitrogen, oxygen in ring, other oxygen, phosphorus, sulfur, halogen exclude fluorine, and the number of rings) for every ligand. The ligand feature profiles between each ligand pair were used to calculate the compound similarity also by the weighted Jaccard coefficient. Using the compound similarity scores (0−1), the protease−ligand complexes were clustered two-way hierarchically using the Euclidean distance method using the server Morpheus (https://software.broadinstitute.org/morpheus/) to generate clustering profiles and distinct clustered groups.

### Antibacterial growth measurements and determination of IC_50_ values

#### Bacterial Growth Rate Determination

E. coli cultures were grown in a M9 minimal medium supplemented with glucose. This was done at a constant temperature of 37°C. After allowing the bacteria to grow overnight, we assessed the density of the bacterial cultures using optical density (OD) measurements at 600 nm. The cultures were then adjusted (or "normalized") to have an OD of 0.1(Palmer, Toprak et al. 2015, Rodrigues and Shakhnovich 2019). This ensures that the starting concentration of bacteria is consistent across different samples. We repeated this normalization process after a further growth period of 5–6 hours.

#### Compound Incubation

Following the growth and normalization, the bacterial cultures were exposed to various concentrations of a known antibiotic (the "positive control"), TMP, and other potential antibiotic compounds ("hit compounds"). These compounds were added to the cultures in 96-well plates, a format that allows for simultaneously testing multiple samples. Roughly, each compound was diluted to one-fifth of its original concentration before being added to the cultures as the highest concentration on the first columns of the 96 well plate followed by serial dilutions in the next 10 columns and keeping the 12th column for the untreated control cell set.

#### Growth Measurement

The plates containing the bacterial cultures and the compounds were then incubated at 37°C. The plates were subjected to orbital shaking, a gentle circular agitation to ensure uniform growth and exposure to the compounds. Every 15 to 30 minutes over a 15-hour period, the optical density (OD) at 600 nm of each culture was measured(Rodrigues and Shakhnovich 2019). This provides a continuous record of bacterial growth in the presence of the compounds. Quantifying Growth: To analyze the growth data, the area under the curve (AUC) of the growth curve (which plots OD against time) was calculated. This AUC provides a single value that represents the total bacterial growth over the 15-hour period. The method for doing this was based on a previously published study by Palmer et al. in 2015.

#### Normalization and IC_50_ Determination

The growth measurements, represented by the AUC values, were then normalized based on the growth of the control samples (i.e., the bacterial cultures not exposed to any hit compounds). This normalization process ensures that any effects observed can be attributed to the hit compounds and no other factors. After normalization, the growth data was plotted against the compound concentrations, and a logistic equation was used to fit the data. From this fitting, the IC_50_ values were determined. The IC50 value represents the concentration of the compound required to inhibit bacterial growth by 50% and is a commonly used metric to assess the potency of an antibiotic. The final reported IC50 values are average values based on multiple (at least three) repeat experiments, and any variability in the measurements is indicated as standard errors.

### Recovery experiments

Wild-type Escherichia coli BW25113 cells underwent cultivation in M9 media, both with and without the addition of potential compounds, enriched with 0.8 g/L glucose and a folate supplement. This folate supplement was composed of 38 μg/mL of both glycine and serine, 75.5 μg/mL of methionine, 1 μg/mL of pantothenate, 20 μg/mL of adenosine, and 50 μg/mL of thymidine. If a compound disrupts metabolic processes within the folate pathway, the addition of the folate mix can alleviate this effect. Consequently, if the folate mix mitigates the disruption caused by a compound, it implies that the compound interferes with a protein in the folate pathway. Considering folB and folX are proteins within this pathway, their recovery through this method indicates that the test compound likely targets these specific proteins. E. coli BW25113 cells, both untreated and treated with candidate compounds, were grown in M9 minimal media supplemented with glucose and the folate mixture at a concentration of 0.8 g/L. We then assessed the growth patterns of these cells across a range of compound concentrations(Chowdhury, Zielinski et al. 2023).

### Overexpression experiments

WT Escherichia coli BL21-RIL-X cells were transformed with plasmids expressing folB and folX genes under pFLAG promoter. Expression was induced with 0.01mM IPTG to minimize the chances of overexpression-induced toxicity. In a 96-well format, both IPTG-induced and uninduced E. coli cells were grown against a concentration gradient of test compounds(Chowdhury, Zielinski et al. 2023). Growth rates were calculated as per the method described above.

### Experimental Evolution Under Arbutin, Metoclopramide, and Risedronic Acid

We performed experimental evolution of *E. coli* BW27783 under sustained exposure to three compounds: the dual folB/folX inhibitor Arbutin, the folB inhibitor Metoclopramide, and the folX inhibitor Risedronic acid. To begin each trajectory, we streaked BW27783 on M9-glucose agar and selected single colonies. We inoculated each colony into 5 mL M9 minimal medium (5X M9 salts diluted to 1X, 0.8 g L⁻¹ glucose, 1 mM MgSO₄, 0.1 mM CaCl₂, and 5 µg mL⁻¹ thiamine) and incubated cultures overnight at 37 °C with shaking at 250 r.p.m(Rodrigues and Shakhnovich 2019, Zhang, Chowdhury et al. 2021). The following day, we diluted each culture into pre-warmed M9 medium to an OD₆₀₀ of ∼0.03 and allowed the cells to resume exponential growth. All evolution experiments were carried out in M9 to ensure a defined metabolic background and to sensitively resolve perturbations to the folate pathway. For each compound, we initiated three independent evolution trajectories. Once cultures reached stable exponential growth, we added the corresponding compound at 0.75 times the IC₅₀ measured for the ancestral strain. This initial concentration imposed strong, sublethal stress that reliably slowed growth while allowing adaptive variants to arise and expand.

We propagated each trajectory by transferring 1-2% (v/v) of the culture every 24 h into fresh M9 medium supplemented with the appropriate drug. Before each passage, we monitored OD₆₀₀ time courses to estimate the growth rate. When a population reached a growth rate comparable to untreated cells, we increased the drug concentration for the next passage. When growth slowed or remained unchanged, we maintained the same drug level. This incremental adjustment maintained continuous selection without risking extinction. We froze aliquots from each trajectory in 25% glycerol. At defined timepoints, we measured full dose–response curves for Arbutin, Metoclopramide, and Risedronic acid using the same concentration ranges applied to the ancestral strain. These measurements enabled us to quantify shifts in IC₅₀ values and follow the progressive accumulation of resistance across the 41-day evolution experiment. After 41 days of serial passaging, we plated each final population on M9-glucose agar and selected individual colonies. We grew two clones from each trajectory in drug-free M9 medium and performed complete inhibition assays to determine clone-specific changes in drug sensitivity relative to the ancestral strain.

### Whole-genome sequencing

We performed whole-genome sequencing on single colonies isolated from the final day of each evolution trajectory. We extracted genomic DNA from each clone and submitted the samples to Novogene for Illumina MiSeq sequencing using a 2 × 150 bp paired-end configuration. We processed the raw reads using the breseq pipeline(Zhang, Chowdhury et al. 2021) with default parameters and aligned them to the *E. coli* BW27783 reference genome (GenBank accession CP009273.1). This analysis allowed us to identify single-nucleotide substitutions, small indels, and structural variants associated with evolved responses to Arbutin, Metoclopramide, and Risedronic acid.

### Enrichment analysis of gene categories within duplicated regions

We performed enrichment analysis to determine whether specific functional gene categories were overrepresented within duplicated genomic regions compared to random expectation. Enrichment analysis tests whether the observed frequency of a given category exceeds what would occur by chance under a defined null model, allowing us to distinguish selective patterns from stochastic variation. The null model posits that the observed locus of genome rearrangement containing duplicated genes is a result of stochastic duplication rather than selected to provide a boost in copy number of genes essential for resistance.

We applied this approach to assess whether gene duplications arising during evolution preferentially target specific functional modules, which would indicate adaptive selection rather than random genome rearrangement. We identified duplicated regions from whole-genome sequencing data using read-depth profiles generated by breseq, where increased coverage relative to the genomic baseline indicates copy number amplification(Deatherage and Barrick 2014).

We annotated genes within each duplicated segment into functional categories, including metabolic, regulatory, stress response, efflux, and folate-related pathways. The folate-related category included both canonical folate biosynthesis genes (fol) and additional genes linked to one-carbon and nucleotide metabolism, capturing both direct pathway components and compensatory metabolic responses.

To assess enrichment, we generated 10,000 random genomic segments of identical length to each duplication across the circular *E. coli* genome. We matched each random segment to the observed duplication in length to control for segment-size–dependent biases in gene category representation. For each segment, we calculated category proportions to construct a null distribution. We computed empirical p-values for enrichment as the fraction of random segments with equal or greater category representation and corrected for multiple testing using the Benjamini-Hochberg procedure (adjusted p < 0.05).

This framework allows us to distinguish selective enrichment of functional modules from stochastic genome rearrangements(Good 2005).

### Mutation analysis and functional classification

We identified genetic variants from whole-genome sequencing data using the breseq pipeline(Deatherage and Barrick 2014) with default parameters. We aligned paired-end reads (2 × 150 bp) to the *E. coli* BW27783 reference genome (GenBank accession CP009273.1) and detected single-nucleotide substitutions, small insertions and deletions (indels), and structural variants based on deviations from the reference sequence. We applied quality filters based on read depth, base quality, and strand consistency to retain high-confidence mutations. We manually inspected sequence alignments to confirm variant positions and exclude potential sequencing artefacts.

We annotated all validated mutations using the BW27783 genome annotation to assign gene identity and genomic context. To analyse adaptive patterns at a functional level, we grouped mutated genes into gene families based on shared functional annotations and known biological roles. These categories included metabolic pathways, regulatory systems, DNA repair, transport, stress response, and mobile genetic elements. We compiled mutation profiles for each independently evolved trajectory and compared the recurrence of genes and gene families across conditions.

We quantified recurrence by identifying genes and gene families that appeared in multiple trajectories within the same condition. We then assessed convergence at both the gene level and the functional (gene family) level to distinguish between specific mutational reuse and broader pathway-level adaptation.

This framework allowed us to characterize how different selective pressures shape the distribution, recurrence, and functional organization of mutations, and to relate these patterns to differences in evolutionary constraint and fitness landscapes across conditions.

## Acknowledgements

Authors acknowledge the funding support NIH R35GM139571 to E.S. Yun-Ti Chen is a fellow in the MIT-Novo Nordisk Artificial Intelligence Postdoctoral Fellows Program, which is supported by funding from Novo Nordisk A/S.

## Author contributions

ES and SC conceptualized the overall project outline. SC and YT designed the computational workplan and YT and CH performed the computational analysis. SC and DS designed the experimental work plans. DS and SC performed the experiments and DS performed the evolution studies and whole genome sequencing. SC, DS and AM performed the in vitro biophysical assays. DS and YT collated the data, performed the analyses, prepared figures and wrote the manuscript. DS, YT, AM, SC and ES reviewed the manuscript.

## Supplementary Information

**Supplementary Figure 1.**
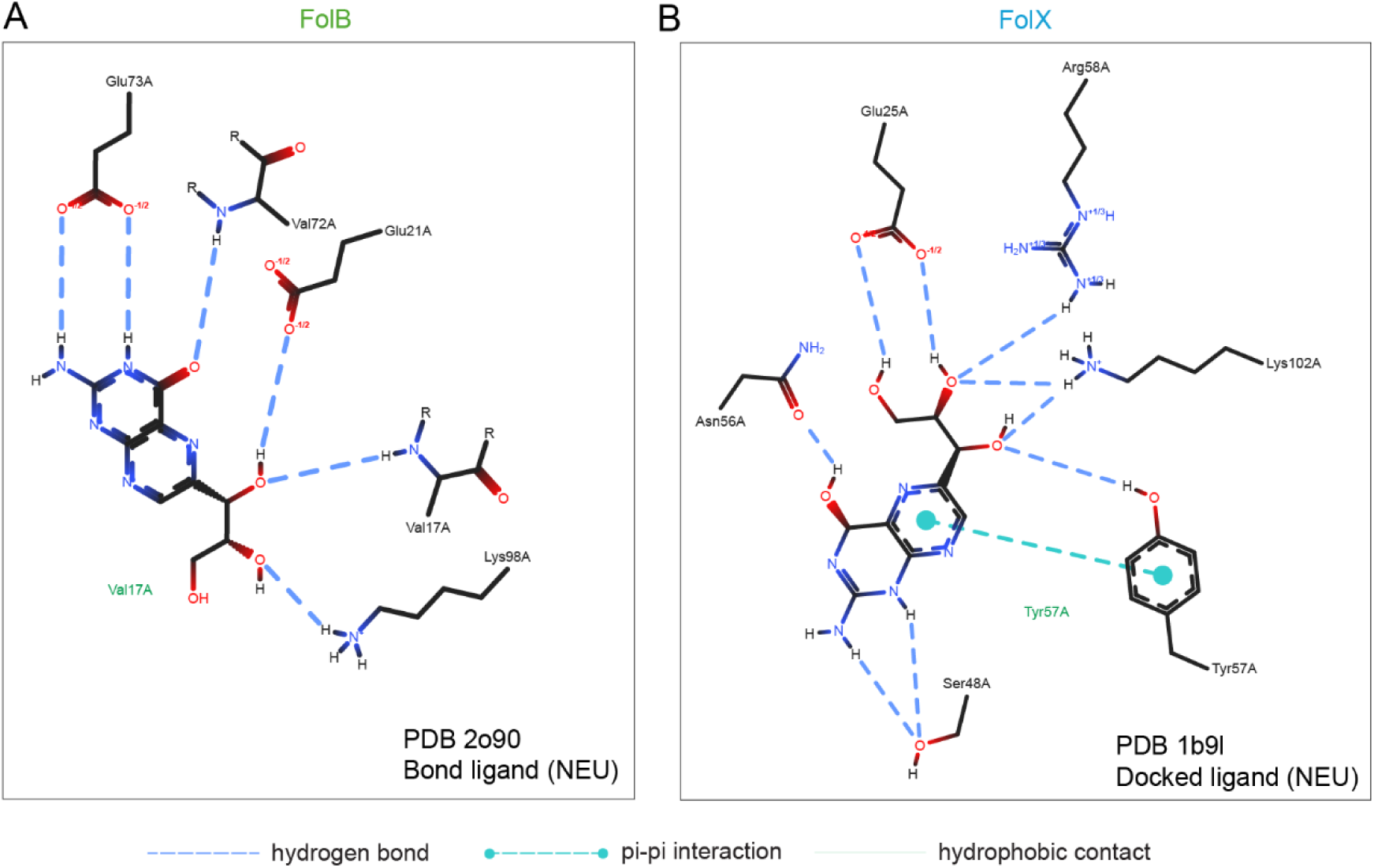
Structural basis of substrate recognition in FolB and FolX. (A) Ligand interaction map of FolB (PDB: 2O90) bound to its native substrate (NEU). The binding pocket is defined by a network of hydrogen bonds involving residues such as Glu73, Val72, Glu21, and Lys98, which stabilize the pterin scaffold and orient the substrate for catalysis. (B) Ligand interaction map of FolX (PDB: 1B9I) with docked NEU. The binding pocket displays a distinct yet partially overlapping interaction network involving residues such as Glu25, Arg58, Lys102, Asn56, Ser48, and Tyr57. In addition to hydrogen bonding, π–π interactions and hydrophobic contacts contribute to ligand stabilization. Dashed lines indicate hydrogen bonds, dotted lines indicate π–π interactions, and grey arcs represent hydrophobic contacts. Together, these interaction patterns highlight conserved physicochemical features of the binding pockets despite sequence divergence.

**Supplementary Figure 2.**
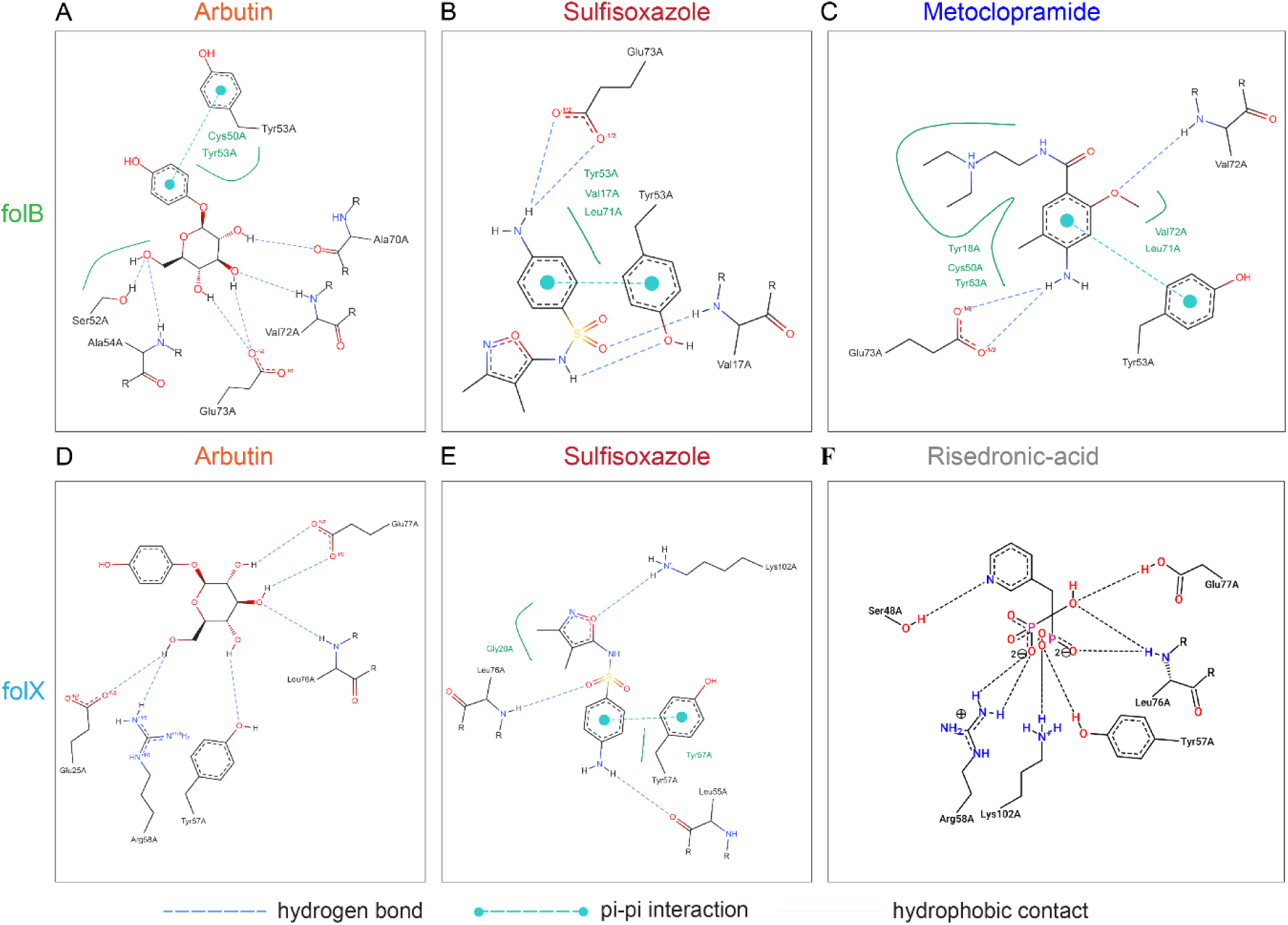
Interaction profiles of representative inhibitors in FolB and FolX binding sites. (A–C) Interaction maps of representative compounds docked in the FolB binding pocket: Arbutin (A), Sulfisoxazole (B), and Metoclopramide (C). These compounds engage conserved residues through combinations of hydrogen bonding and hydrophobic interactions, with aromatic groups positioned to interact with Tyr53 and surrounding hydrophobic residues. (D–F) Interaction maps of representative compounds docked in the FolX binding pocket: Arbutin (D), Sulfisoxazole (E), and Risedronic acid (F). Binding involves a combination of polar contacts with residues such as Glu25, Arg58, and Lys102, along with aromatic stacking interactions near Tyr57, reflecting adaptation to the FolX pocket architecture. Hydrogen bonds are shown as dashed blue lines, π–π interactions as cyan connectors, and hydrophobic contacts as green arcs. These interaction patterns illustrate how chemically diverse compounds achieve binding through conserved interaction hotspots across both enzymes.

**Supplementary Figure 3.**
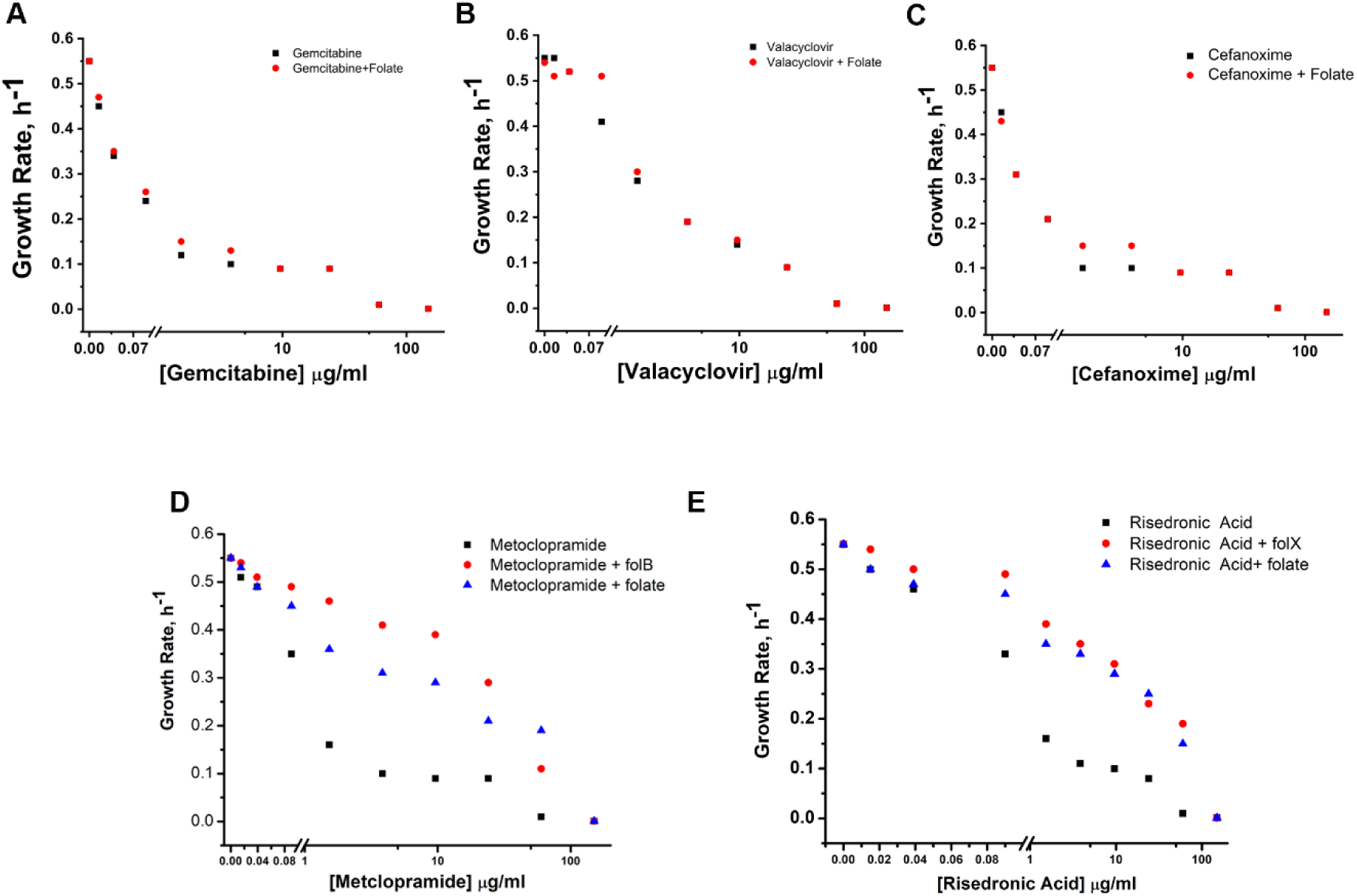
Growth rates of bacteria treated with increasing concentrations of (B) Arbutin, (A) Gemcitabine, (B) Valacyclovir, (C) Cefanoxime (D) Metoclopramide, and (E) Risedronic acid measured with (red) and without (black) exogenous folate. Folate restores growth in (D) Metoclopramide, and (E) Risedronic acid treated cells, while no rescue pattern was observed for other inhibitors. Growth rates under Metoclopramide and Risedronic acid exposure in strains overexpressing folB and folX showed rescue trends

**Supplementary Figure 4.**
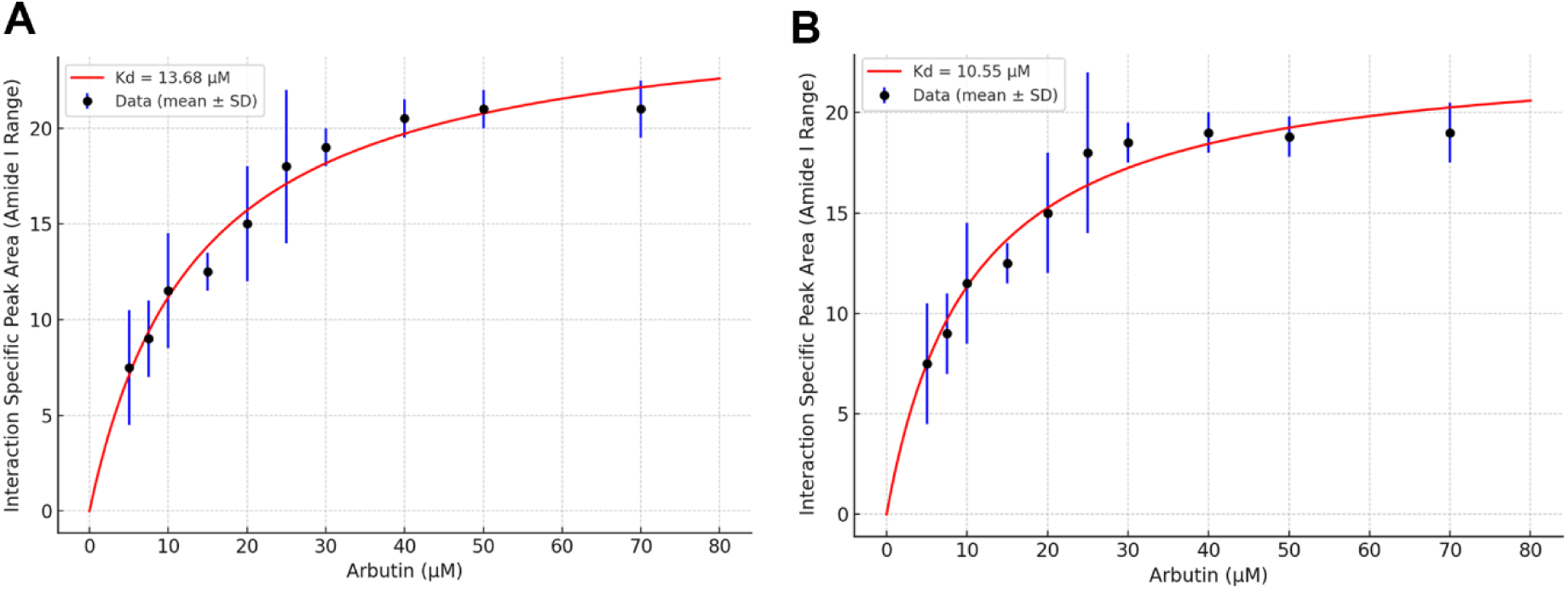
Raman spectroscopy based binding experiments show arbutin co-targets folB and folX. (A-B) Binding interaction profiles of arbutin with folB and folX. Hyperbolic binding fits show concentration-dependent interaction-specific peak area (amide-I range) for (A) folB and (B) folX, with extracted dissociation constants indicating measurable association between arbutin and each target. Data points represent mean ± SD. We normalized and scaled Raman intensities (including log transformation where appropriate) to enable comparison across proteins and conditions. Because the signal reflects ligand-induced structural perturbations, the derived K_d_ values represent effective interaction parameters rather than direct thermodynamic binding affinities and may differ from values obtained using thermodynamic methods (e.g., ITC).

**Supplementary Figure 5.**
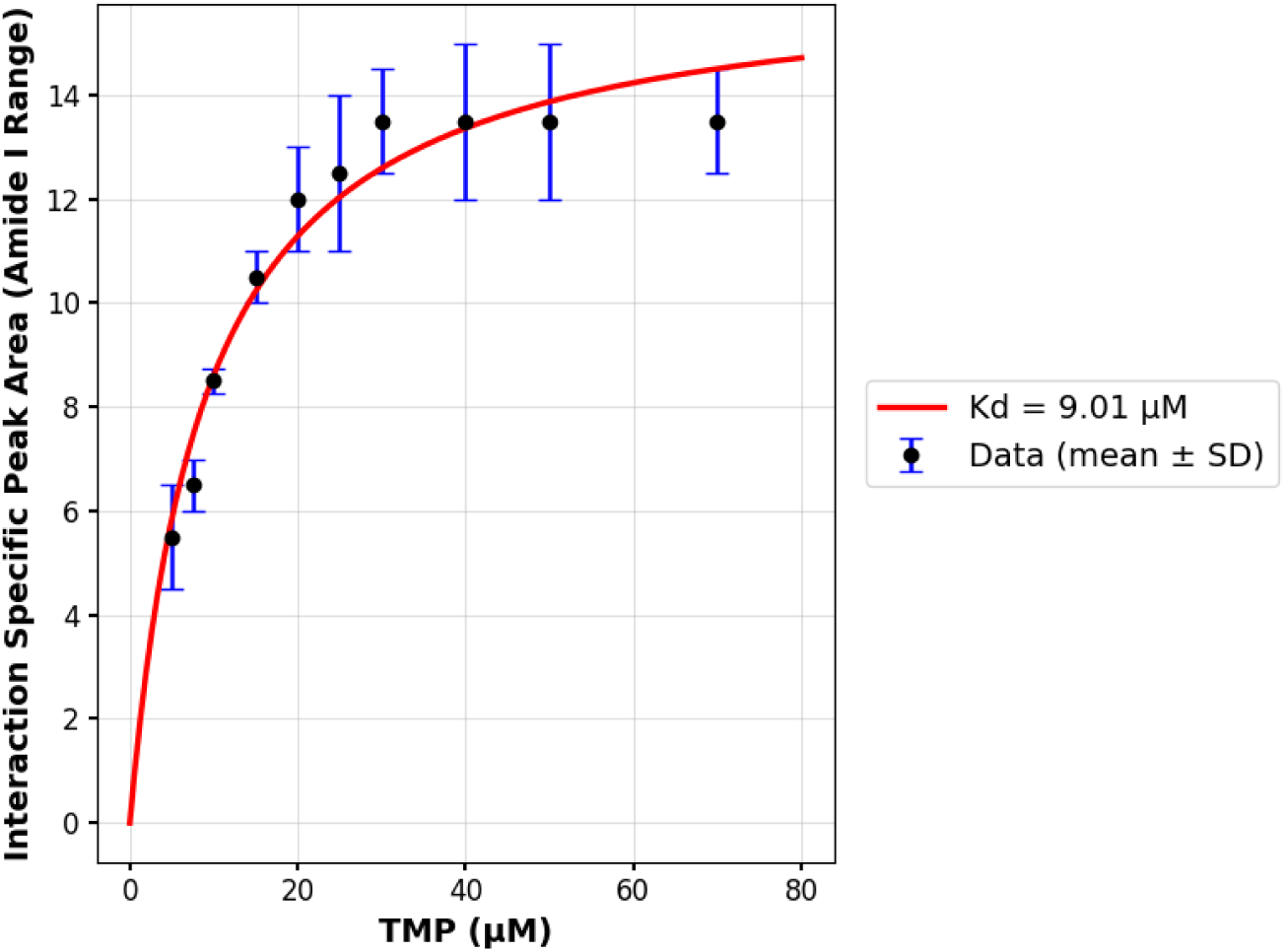
Binding interaction profiles of TMP (Trimethoprim) with folA (DHFR). Hyperbolic binding fits show concentration-dependent interaction-specific peak area (amide-I range). Data points represent mean ± SD.

**Supplementary Figure 6.**
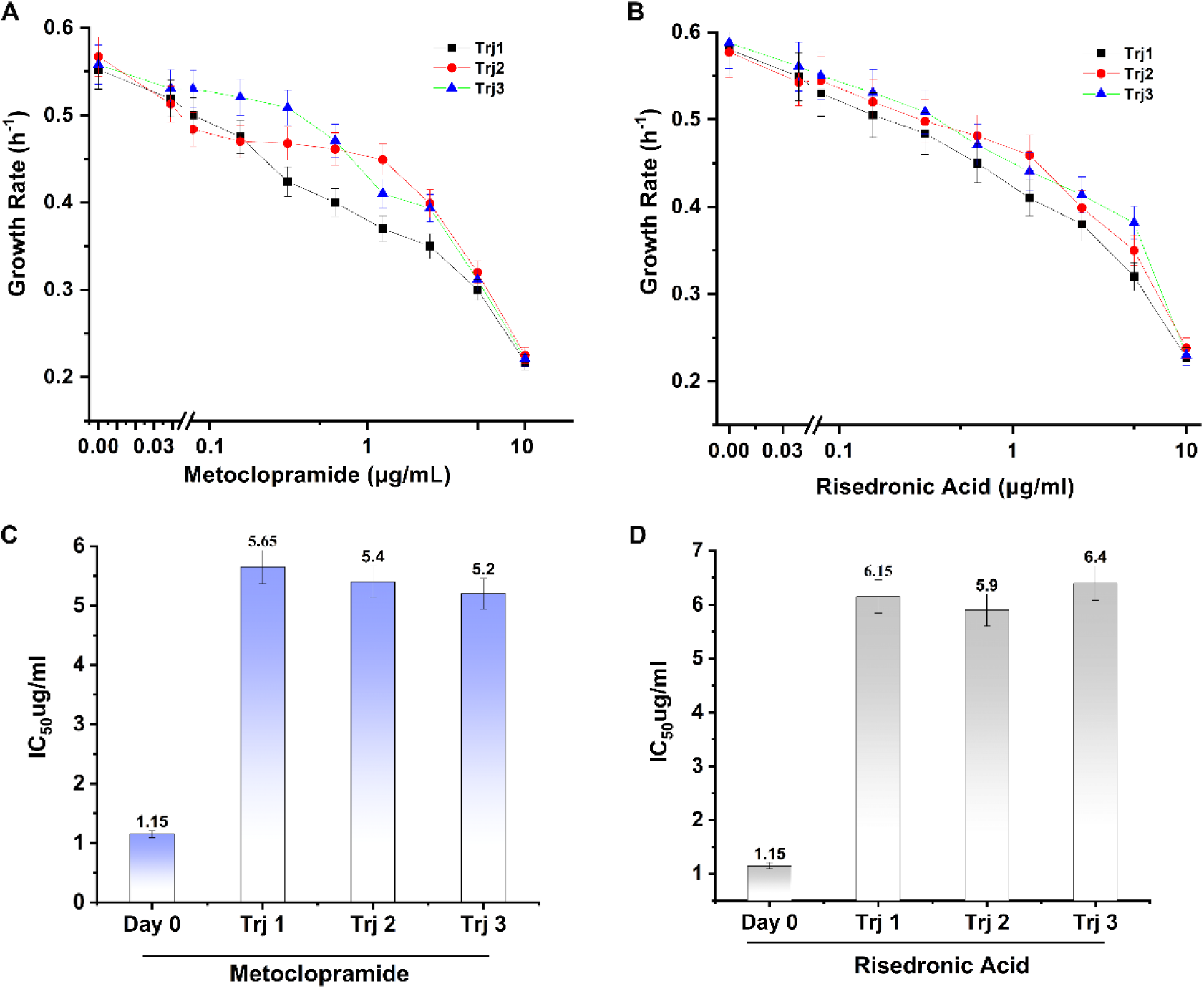
Evolution of resistance under single-target inhibition of FolB and FolX. (A) Growth-rate measurements for populations evolved under Metoclopramide (FolB inhibitor). Three independent trajectories show reduced growth rates with increasing Metoclopramide concentration, while maintaining higher growth at intermediate concentrations relative to the ancestral strain. Points represent mean values; error bars indicate standard deviation. (B) Growth-rate measurements for populations evolved under Risedronic acid (FolX inhibitor). Three independent trajectories display a progressive reduction in growth rate with increasing drug concentration. Points represent mean values; error bars indicate standard deviation. (C) IC_50_ values for Metoclopramide-evolved populations. The ancestral IC_50_ (1.15 µg/mL) is shown on the left. Evolved populations exhibit elevated IC_50_ values ranging from ∼5.2 to 5.7 µg/mL across independent trajectories. (D) IC_50_ values for Risedronic acid–evolved populations. The ancestral IC_50_ (1.15 µg/mL) is shown on the left. Evolved populations show increased IC_50_ values ranging from ∼5.9 to 6.4 µg/mL across independent trajectories.

**Supplementary Table 1.** 19 tested drugs and their IC_50_ from growth measurements.

| No. | Name | Predicted target | IC<sub>50</sub> (µg/ml) |
| --- | --- | --- | --- |
| 1 | Valganciclovir | Both folB and folX | 1.9 |
| 2 | Arbutin | Both folB and folX | 1.3 |
| 3 | Sulfisoxazole | Both folB and folX | 0.25 |
| 4 | Nitenpyram | Both folB and folX | 2.5 |
| 5 | Cytarabine | Both folB and folX | N/A |
| 6 | Droxidopa | Both folB and folX | N/A |
| 7 | Vidarabine | Both folB and folX | N/A |
| 8 | Gemcitabine | folB | 0.35 |
| 9 | Valaciclovir | folB | 0.9 |
| 10 | Cefmenoxime | folB | 0.2 |
| 11 | Amiloride | folB | N/A |
| 12 | Lodoxamide | folB | N/A |
| 13 | Metoclopramide | folB | N/A |
| 14 | Ribavirin | folB | N/A |
| 15 | Zanamivir | folB | N/A |
| 16 | Aztreonam | folX | 1.2 |
| 17 | Nelarabine | folX | N/A |
| 18 | Nitazoxanide | folX | N/A |
| 19 | Risedronic Acid | folX | N/A |

**Supplementary Table 2.** Functional classification of mutations in the Arbutin-evolved trajectory Trj1.

| Start | End | Variant Type | Gene Name | CDS | Gene_type |
| --- | --- | --- | --- | --- | --- |
| 3825094 | 3825095 | snp | ispD | 4-diphosphocytidyl-2C-methyl-D-erythritol synthase | Metabolic |
| 3808101 | 3808102 | snp | pphB | serine/threonine-specific protein phosphatase 2 | Others |
| 3620491 | 3620491 | deletion | nadB | quinolinate synthase, L-aspartate oxidase (B protein) subunit | Metabolic |
| 3513759 | 3513760 | snp | yfgF | cyclic-di-GMP phosphodiesterase, anaerobic | Regulatory |
| 3507298 | 3508521 | insertion | ppk | polyphosphate kinase, component of RNA degradosome | Metabolic |
| 3375626 | 3375627 | snp | yfdX | uncharacterized protein | Others |
| 3174042 | 3174043 | snp | arnA | fused UDP-L-Ara4N formyltransferase/UDP-GlcA C-4'-decarboxylase | Others |
| 2781716 | 2781897 | insertion | yeeA | putative transporter, FUSC family inner membraneprotein | Cell envelope |
| 2579380 | 2579380 | deletion | yobB | C-N hydrolase family protein | Others |
| 2542679 | 2542680 | snp | sdaA | L-serine dehydratase 1 | Metabolic |
| 2280353 | 2281130 | insertion | uidA | beta-D-glucuronidase | Metabolic |
| 2226965 | 2226966 | snp | ynfA | UPF0060 family inner membrane protein | Cell envelope |
| 1900802 | 1900802 | deletion | ydaM | diguanylate cyclase, csgD regulator | Regulatory |
| 1710335 | 1710335 | deletion | ychF | putative GTP-binding protein; catalase inhibitor protein | Regulatory |
| 1362767 | 1362767 | deletion | ompF | outer membrane porin 1a (Ia;b;F) | Cell envelope |
| 1039807 | 1039808 | snp | rhsO | pseudogene, Rhs family | Others |
| 773309 | 773310 | snp | ybcF | putative carbonate kinase | Metabolic |
| 459495 | 459496 | snp | yahO | periplasmic protein | Cell envelope |
| 459494 | 459495 | snp | yahO | periplasmic protein | Cell envelope |
| 459493 | 459494 | snp | yahO | periplasmic protein | Cell envelope |
| 323274 | 323275 | insertion | gpt | xanthine phosphoribosyltransferase; xanthine-guanine phosphoribosyltransferase | Metabolic |
| 4419216 | 4419217 | snp | nusA | transcription termination/antitermination L factor | Regulatory |
| 4398233 | 4398234 | snp | yhbT | SCP-2 sterol transfer family protein | Others |
| 4355170 | 4355171 | snp | tdcR | L-threonine dehydratase operon activator protein | Regulatory |
| 4352295 | 4352296 | snp | tdcB | L-threonine dehydratase, catabolic | Metabolic |
| 4220150 | 4220151 | snp | mqsA | antitoxin for MqsR toxin; transcriptional repressor | Efflux and Stress response |
| 4220149 | 4220149 | deletion | mqsA | antitoxin for MqsR toxin; transcriptional repressor | Efflux and Stress response |
| 4041695 | 4041696 | snp | recJ | ssDNA exonuclease, 5' --> 3'-specific | Regulatory |
| 3931049 | 3931050 | snp | recD | exonuclease V (RecBCD complex), alpha chain | Regulatory |
| 3914020 | 3914021 | snp | rlmM | 23S rRNA C2498 2'-O-ribose methyltransferase, SAM-dependent | Regulatory |
| 3899860 | 3899861 | snp | sdaB | L-serine dehydratase 2 | Metabolic |

**Supplementary Table 3.** Functional classification of mutations in the Arbutin-evolved trajectory Trj2.

| Start | End | Variant Type | Gene Name | CDS | Gene_type |
| --- | --- | --- | --- | --- | --- |
| 3825094 | 3825095 | snp | metZ | Unknown | Others |
| 3808101 | 3808102 | snp | fucP | L-fucose transporter | Others |
| 3620491 | 3620491 | deletion | csiD | carbon starvation protein | Efflux and Stress response |
| 3513759 | 3513760 | snp | yfhH | putative DNA-binding transcriptional regulator | Regulatory |
| 3507298 | 3508521 | insertion | purL | phosphoribosylformyl-glycineamide synthetase | Metabolic |
| 3375626 | 3375627 | snp | acrD | aminoglycoside | Regulatory |
| 3174042 | 3174043 | snp | folC | bifunctional folylpolyglutamate synthase/ dihydrofolate synthase | Folate pathway |
| 2781716 | 2781897 | insertion | wzc | colanic acid production tyrosine-protein kinase;autokinase; Ugd phosphorylase | Metabolic |
| 2579380 | 2579380 | deletion | cheA | sensory histidine kinase/signal sensing protein | Regulatory |
| 2542679 | 2542680 | snp | ruvB | ATP-dependent DNA helicase, component of RuvABC resolvasome | Metabolic |
| 2280353 | 2281130 | insertion | sodB | superoxide dismutase, Fe | Efflux and Stress response |
| 2226965 | 2226966 | snp | uidB | glucuronide transporter | Others |
| 1900802 | 1900802 | deletion | ydbH | putative membrane-anchored protein | Cell envelope |
| 1710335 | 1710335 | deletion | rssB | PcnB-degradosome interaction factor; response regulator | Regulatory |
| 1362767 | 1362767 | deletion | pqiA | paraquat-inducible, SoxRS-regulated inner membrane protein | Cell envelope |
| 1039807 | 1039808 | snp | sdhA | succinate dehydrogenase, flavoprotein subunit | Metabolic |
| 773309 | 773310 | snp | sfmZ | response regulator family protein | Regulatory |
| 459495 | 459496 | snp | prpE | propionate--CoA ligase | Metabolic |
| 459494 | 459495 | snp | prpE | propionate--CoA ligase | Metabolic |
| 459493 | 459494 | snp | prpE | propionate--CoA ligase | Metabolic |
| 323274 | 323275 | insertion | ykfI | CP4-6 prophage; toxin of the YkfI-YafW toxin-antitoxin system | Efflux and Stress response |
| 4419216 | 4419217 | snp | accB | acetyl CoA carboxylase, BCCP subunit | Metabolic |
| 4398233 | 4398234 | snp | aaeA | p-hydroxybenzoic acid efflux system component | Efflux and Stress response |
| 4355170 | 4355171 | snp | gltB | glutamate synthase, large subunit | Metabolic |
| 4352295 | 4352296 | snp | arcB | aerobic respiration control sensor histidine protein kinase, cognate to two-component response regulators ArcA and RssB | Regulatory |
| 4220150 | 4220151 | snp | rlmG | 23S rRNA m(2)G1835 methyltransferase, SAM-dependent | Metabolic |
| 4220149 | 4220149 | deletion | rlmG | 23S rRNA m(2)G1835 methyltransferase, SAM-dependent | Metabolic |
| 4041695 | 4041696 | snp | pppA | bifunctional prepilin leader peptidase/ methylase | Metabolic |
| 3931049 | 3931050 | snp | uacT | uric acid permease | Others |

**Supplementary Table 4.** Functional classification of mutations in the Arbutin-evolved trajectory Trj3.

| Start | End | Variant Type | Gene Name | CDS | Gene_type |
| --- | --- | --- | --- | --- | --- |
| 3824592 | 3824593 | snp | xdhD | putative hypoxanthine oxidase, molybdopterin-binding/Fe-S binding | Folate pathway |
| 3807918 | 3807919 | snp | hyuA | D-stereospecific phenylhydantoinase | Metabolic |
| 3620308 | 3620308 | deletion | pphB | serine/threonine-specific protein phosphatase 2 | Metabolic |
| 3513759 | 3513760 | snp | yfjW | CP4-57 prophage; putative inner membrane protein | Recombination |
| 3507298 | 3508521 | insertion | yfjQ | CP4-57 prophage; uncharacterized protein | Recombination |
| 3375626 | 3375627 | snp | hscB | HscA co-chaperone, J domain-containing protein Hsc56; IscU-specific chaperone HscAB | Others |
| 3174042 | 3174043 | snp | ypdA | sensor kinase regulating yhjX; pyruvate-responsive YpdAB two-component system | Regulatory |
| 2781716 | 2781897 | insertion | metG | methionyl-tRNA synthetase | Metabolic |
| 2579380 | 2579380 | deletion | hchA | glyoxalase III and Hsp31 molecular chaperone | Others |
| 2542679 | 2542680 | snp | amyA | cytoplasmic alpha-amylase | Metabolic |
| 2280353 | 2281130 | insertion | btuC | vitamin B12 ABC transporter permease | Others |
| 2226965 | 2226966 | snp | ydhX | putative 4Fe-4S ferredoxin-type protein; FNR, Nar, NarP-regulated protein | Metabolic |
| 1900802 | 1900802 | deletion | trg | methyl-accepting chemotaxis protein III, ribose and galactose sensor receptor | Regulatory |
| 1710335 | 1710335 | deletion | osmB | osmotically and stress inducible lipoprotein | Efflux and Stress response |
| 1362767 | 1362767 | deletion | torR | response regulator in two-component regulatory system with TorS | Regulatory |
| 1039807 | 1039808 | snp | ybhA | pyridoxal phosphate (PLP) phosphatase | Metabolic |
| 773309 | 773310 | snp | nfrB | bacteriophage N4 receptor, inner membrane subunit | Recombination |
| 459495 | 459496 | snp | lacZ | beta-D-galactosidase | Metabolic |
| 459494 | 459495 | snp | lacZ | beta-D-galactosidase | Metabolic |
| 459493 | 459494 | snp | lacZ | beta-D-galactosidase | Metabolic |
| 323274 | 323275 | insertion | yafZ | CP4-6 prophage; conserved protein | Recombination |
| 4419033 | 4419034 | snp | tsgA | putative transporter | Others |
| 4398050 | 4398051 | snp | yheO | putative PAS domain-containing DNA-binding transcriptional regulator | Regulatory |
| 4354987 | 4354988 | snp | rpoA | RNA polymerase, alpha subunit | Regulatory |
| 4352112 | 4352113 | snp | trkA | NAD-binding component of Trk potassium transporter | Others |
| 4219967 | 4219968 | snp | dacB | D-alanyl-D-alanine carboxypeptidase | Metabolic |
| 4219966 | 4219966 | deletion | dacB | D-alanyl-D-alanine carboxypeptidase | Metabolic |
| 4041512 | 4041513 | snp | glnE | fused deadenyltransferase/adenylyltransferase for glutamine synthetase | Metabolic |
| 3930866 | 3930867 | snp | speC | ornithine decarboxylase, constitutive | Metabolic |

**Supplementary Table 5.** Gene family-level similarity of mutations across Arbutin-evolved trajectories.

| Gene family (name-based) | Trj1 genes | Trj2 genes | Trj3 genes | Similarity Pattern |
| --- | --- | --- | --- | --- |
| rec / recombination | recJ, recD | — | — | **Trj1-specific (repair family hit)** |
| ruv / recombination | — | ruvB | — | **Trj2-specific (repair family hit)** |
| DNA repair helicase/exonuclease machinery (rec+ruv as a unit) | recJ, recD | ruvB | – | **Yes (Trj1 ↔ Trj2) at the “recomb/repair gene-family” level** |
| rlm / rRNA methylation | rlmM | rlmG | – | **Yes (Trj1 ↔ Trj2) (same rlm family, different paralogs)** |
| sda / serine deaminase | sdaA, sdaB | – | – | **Trj1-specific (clear within-trajectory family hit)** |
| tdc / threonine/serine catabolism regulon | tdcR, tdcB | – | – | **Trj1-specific (within-trajectory family hit)** |
| uid / β-glucuronidase operon family | uidA | uidB | – | **Yes (Trj1 ↔ Trj2) (same uid family, different genes)** |
| yfj* | – | – | yfjW, yfjQ | **Trj3-specific (within-trajectory family hit)** |
| ygf* | ygfF | ygfK | – | **Yes (Trj1 ↔ Trj2) (same ygf family, different genes)** |
| met* (methionine-related genes; name family only) | – | metZ | metG | **Weak yes (Trj2 ↔ Trj3) (both “met” family but not same function)** |
| pphB (exact gene repeat) | pphB | – | pphB | **Strong yes (Trj1 ↔ Trj3) (only clear exact gene overlap)** |
| Two-component / sensory kinase-regulator family (name-based examples) | – | arcB, cheA | torR | **Yes (Trj2 ↔ Trj3) (same broad “sensor/regulator” gene-family theme, but names differ)** |

**Supplementary Table 6.**
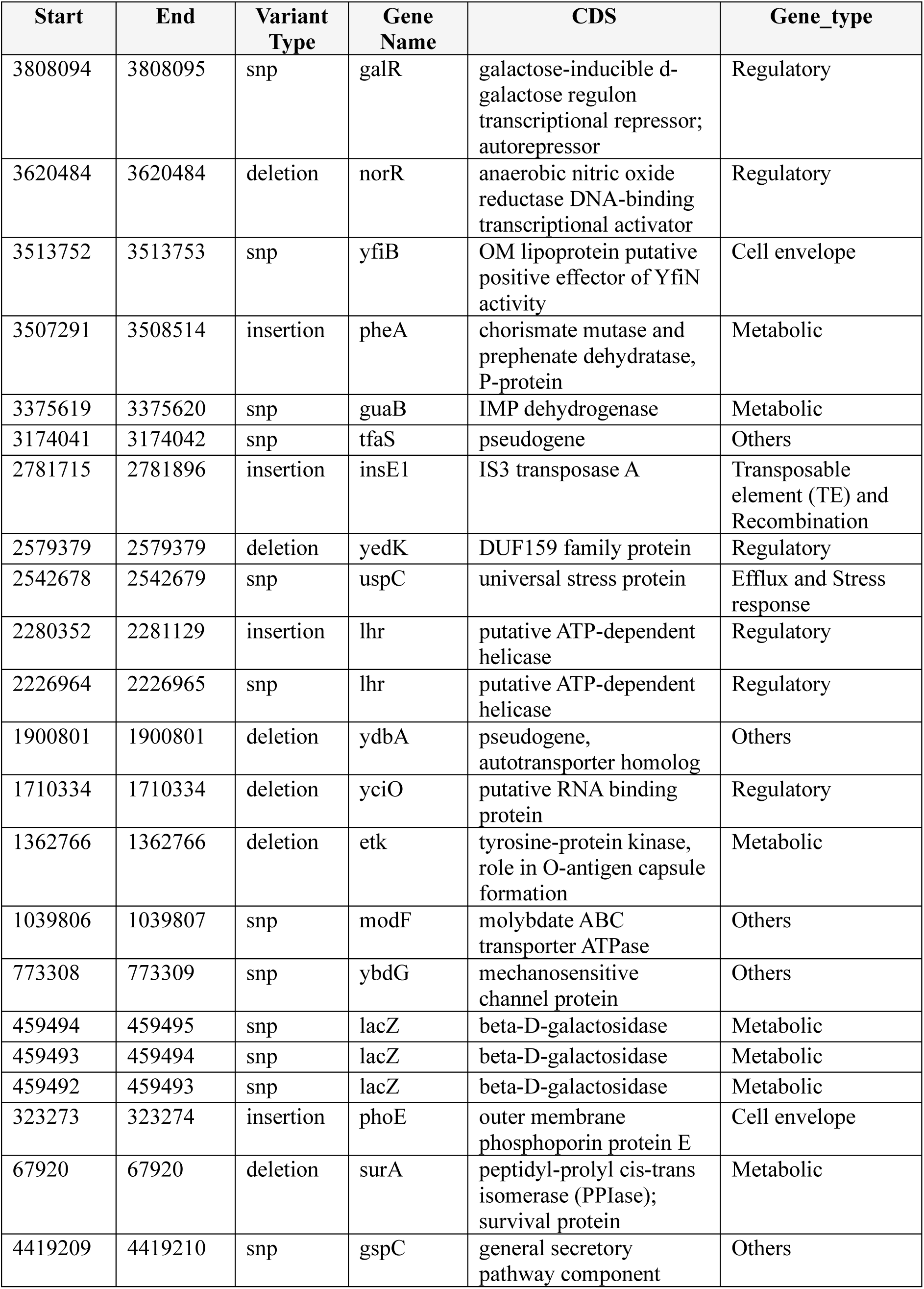
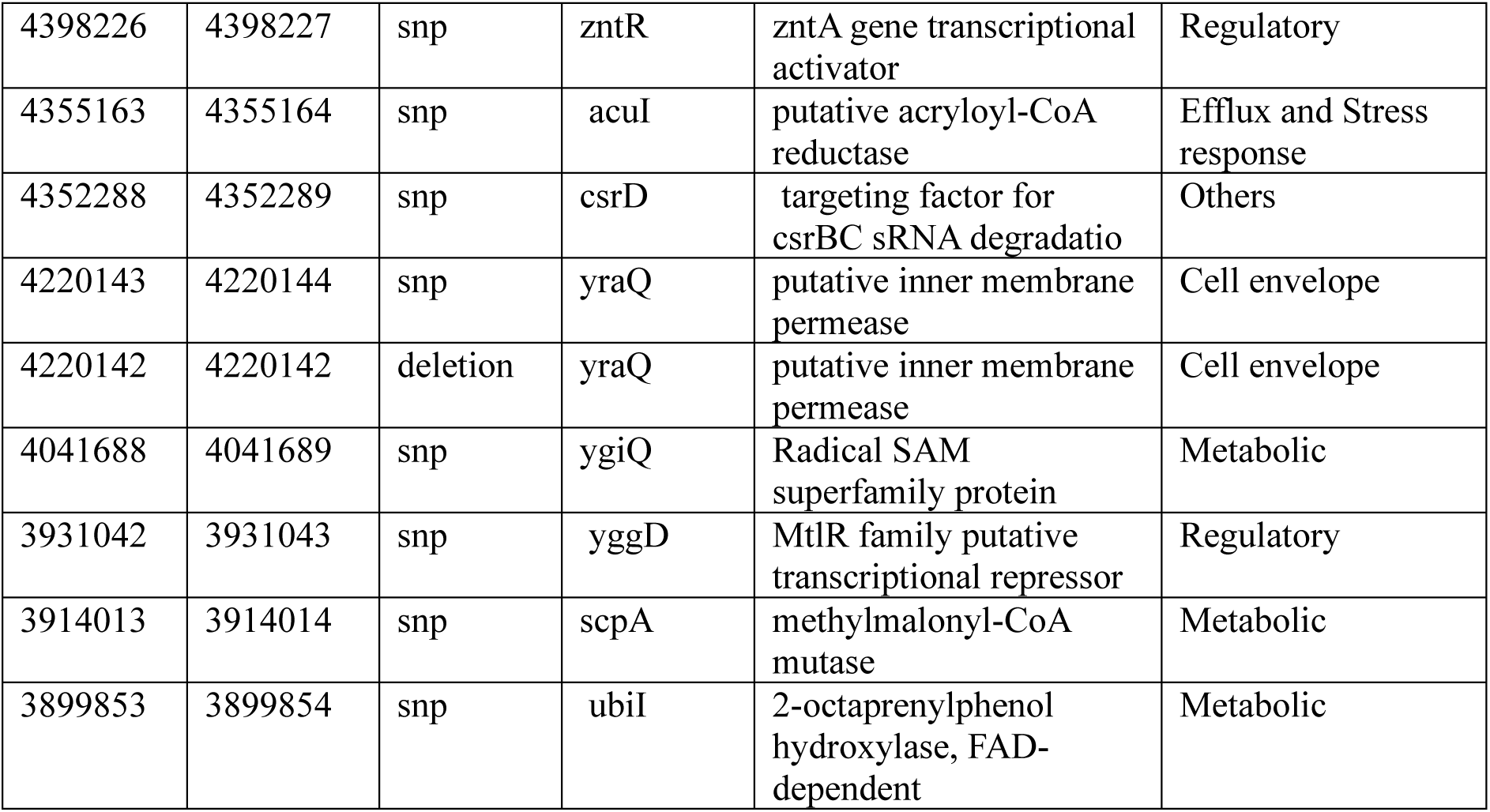
Functional classification of mutations in the Risedronic acid-evolved trajectory Trj1 (folX inhibition).

**Supplementary Table 7.**
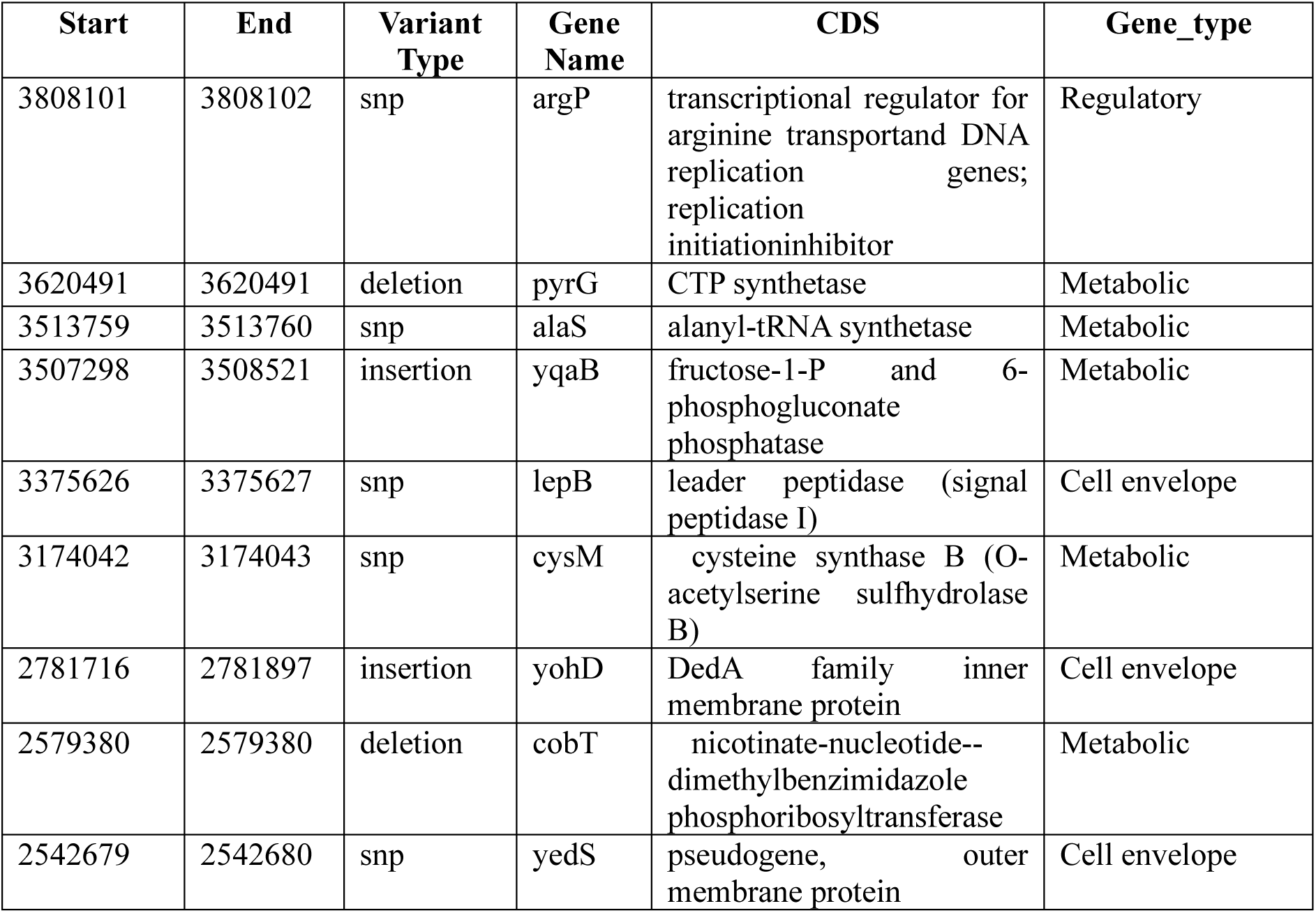
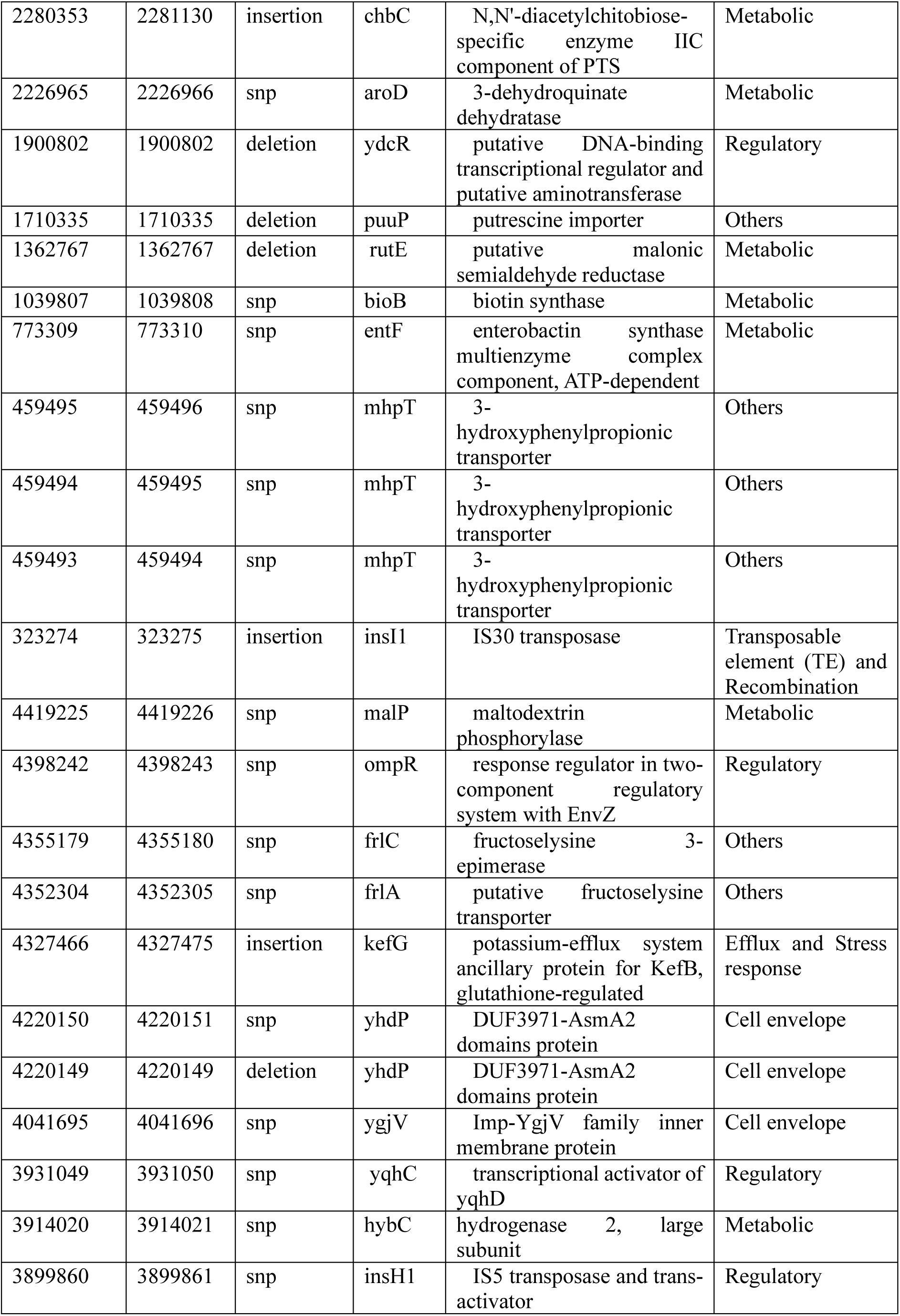
Functional classification of mutations in the Risedronic acid-evolved trajectory Trj2 (folX inhibition).

**Supplementary Table 8.** Functional classification of mutations in the Risedronic acid-evolved trajectory Trj3 (folX inhibition).

| Start | End | Variant Type | Gene Name | CDS | Gene_type |
| --- | --- | --- | --- | --- | --- |
| 3490426 | 3490427 | snp | yfhM | bacterial alpha2-macroglobulin colonization factor ECAM; anti-host protease defense factor | Efflux and Stress response |
| 3678037 | 3678037 | deletion | nrdI | NrdEF cluster assembly flavodoxin | Metabolic |
| 3784768 | 3784769 | snp | casD | CRISP RNA (crRNA) containing Cascade antiviral complex protein | Regulatory |
| 3790007 | 3791230 | insertion | ygcB | Cascade complex anti-viral R-loop helicase-annealase Cas3 | Regulatory |
| 3922901 | 3922902 | snp | yqeH | putative LuxR family transcriptional regulator | Regulatory |
| 4124485 | 4124486 | snp | hybD | maturation protease for hydrogenase 2 | Metabolic |
| 4516631 | 4516812 | insertion | rpsK | 30S ribosomal subunit protein S11 | Regulatory |
| 89892 | 89892 | deletion | araA | L-arabinose isomerase | Metabolic |
| 126592 | 126593 | snp | murD | UDP-N-acetylmuramoyl-L-alanine:D-glutamate ligase | Metabolic |
| 388142 | 388919 | insertion | paoD | moco insertion factor for PaoABC aldehyde oxidoreductase | Others |
| 442306 | 442307 | snp | yahI | carbamate kinase-like protein | Metabolic |
| 768470 | 768470 | deletion | nfrA | bacteriophage N4 receptor, outer membrane subunit | Cell envelope |
| 958937 | 958937 | deletion | pgm | phosphoglucomutase | Metabolic |
| 1306505 | 1306505 | deletion | mukB | chromosome condensin MukBEF, ATPase and DNA-binding subunit | Others |
| 1629464 | 1629465 | snp | minD | inhibitor of FtsZ ring polymerization; chromosome-membrane tethering protein | Cell envelope |
| 1895962 | 1895963 | snp | ttcC | pseudogene, prophage Rac integration site ttcA duplication; Phage or Prophage Related | Transposable element (TE) and Recombination |
| 2209776 | 2209777 | snp | pntA | pyridine nucleotide transhydrogenase, alpha subunit | Metabolic |
| 2209777 | 2209778 | snp | pntA | pyridine nucleotide transhydrogenase, alpha subunit | Metabolic |
| 2209778 | 2209779 | snp | pntA | pyridine nucleotide transhydrogenase, alpha subunit | Metabolic |
| 2345997 | 2345998 | insertion | ydiP | putative DNA-binding transcriptional regulator | Regulatory |
| 2601351 | 2601351 | deletion | cheA | fused chemotactic sensory histidine kinase in two-component regulatory system | Regulatory |
| 2879311 | 2879312 | snp | yehB | putative outer membrane protein | Cell envelope |
| 2900294 | 2900295 | snp | yehM | uncharacterized protein | Others |
| 2943357 | 2943358 | snp | mglA | methyl-galactoside ABC transporter ATPase | Others |
| 2946232 | 2946233 | snp | mglB | methyl-galactoside transporter subunit | Others |
| 3078377 | 3078378 | snp | gyrA | DNA gyrase (type II topoisomerase), subunit A | Metabolic |
| 3078379 | 3078379 | deletion | gyrA | DNA gyrase (type II topoisomerase), subunit A | Metabolic |
| 3256832 | 3256833 | snp | yfdK | CPS-53 (KpLE1) prophage; conserved protein | Transposable element (TE) and Recombination |
| 3367478 | 3367479 | snp | intZ | CPZ-55 prophage; putative phage integrase | Transposable element (TE) and Recombination |
| 3384507 | 3384508 | snp | eutD | phosphate acetyltransferase | Metabolic |
| 3398667 | 3398668 | snp | nudK | GDP-mannose pyrophosphatase | Others |

**Supplementary Table 9.** Gene family-level similarity of mutations across Risedronic acid-evolved trajectories.

| Gene-family (name-based) | Trj1 genes | Trj2 genes | Trj3 genes | Similarity pattern |
| --- | --- | --- | --- | --- |
| ins* (insertion sequence / IS element genes) | insE1 | insI1, insH1 | – | Shared Trj1 ↔ Trj2 (TE/IS family repeatedly hit) |
| yed* | yedK | yedS | – | Shared Trj1 ↔ Trj2 (same yed-family, different members) |
| mgl* | – | – | mglA, mglB | Trj3-specific within-trajectory family hit |
| hyb* (hydrogenase-related genes; name-family) | – | – | hybD, hybC | Trj3-specific within-trajectory family hit |
| frlA / frlC | – | frlA, frlC | – | Trj2-specific within-trajectory family hit |
| lacZ (exact gene) | lacZ | – | – | Trj1-specific |
| regulator “-R” family (name pattern only) | galR, norR, zntR | – | – | Trj1 enriched in “R” regulators (pattern, not paralogs) |
| response regulator / two-component name-family | – | ompR | – | Trj2-specific |
| chemotaxis “che” family | – | – | cheA | Trj3-specific |
| DNA gyrase family | – | – | gyrA | Trj3-specific |
| cell division / partition family (min/muk) | – | – | minD, mukB | Trj3-specific (two separate families but same theme) |
| nucleotide/redox “nud/nrd” families | – | – | nudK, nrdI | Trj3-specific |

**Supplementary Table 10.**
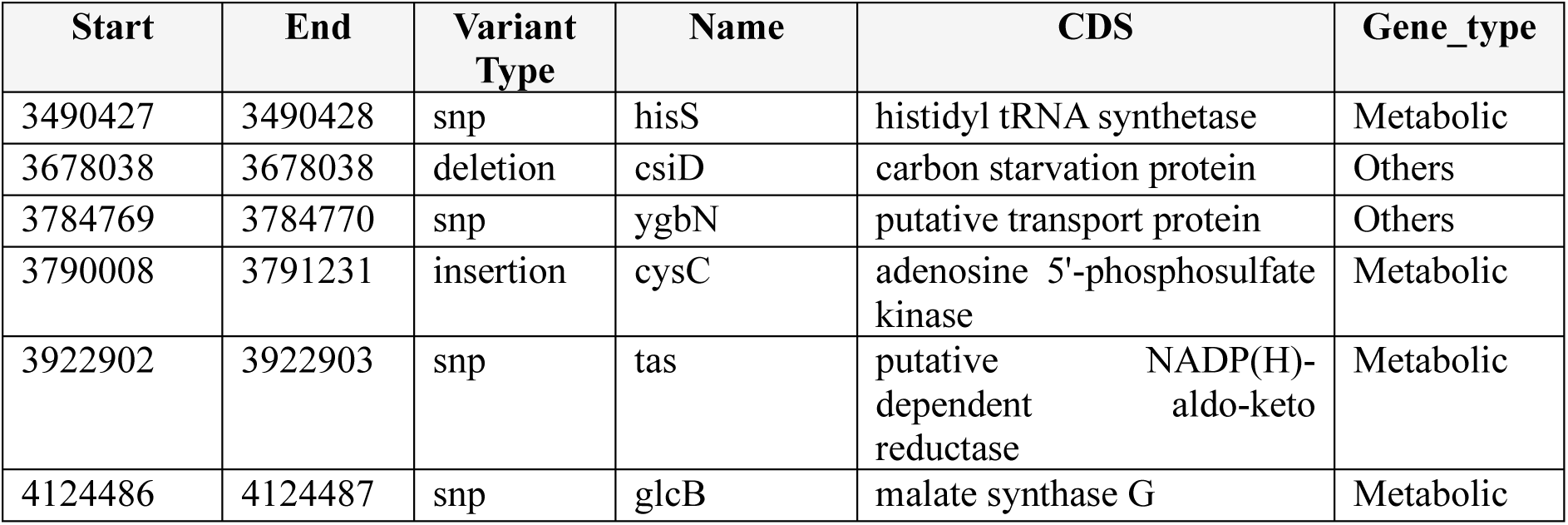
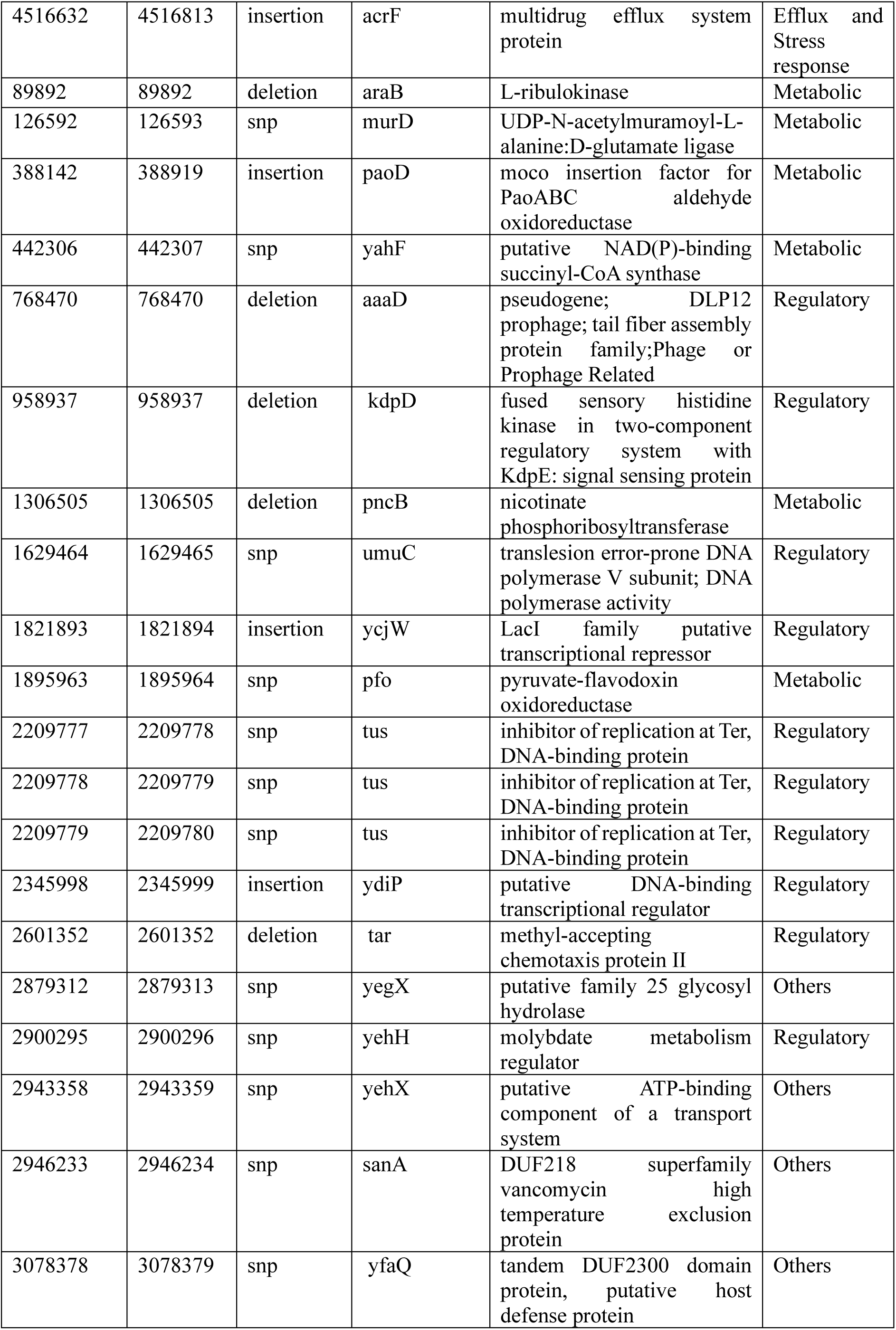
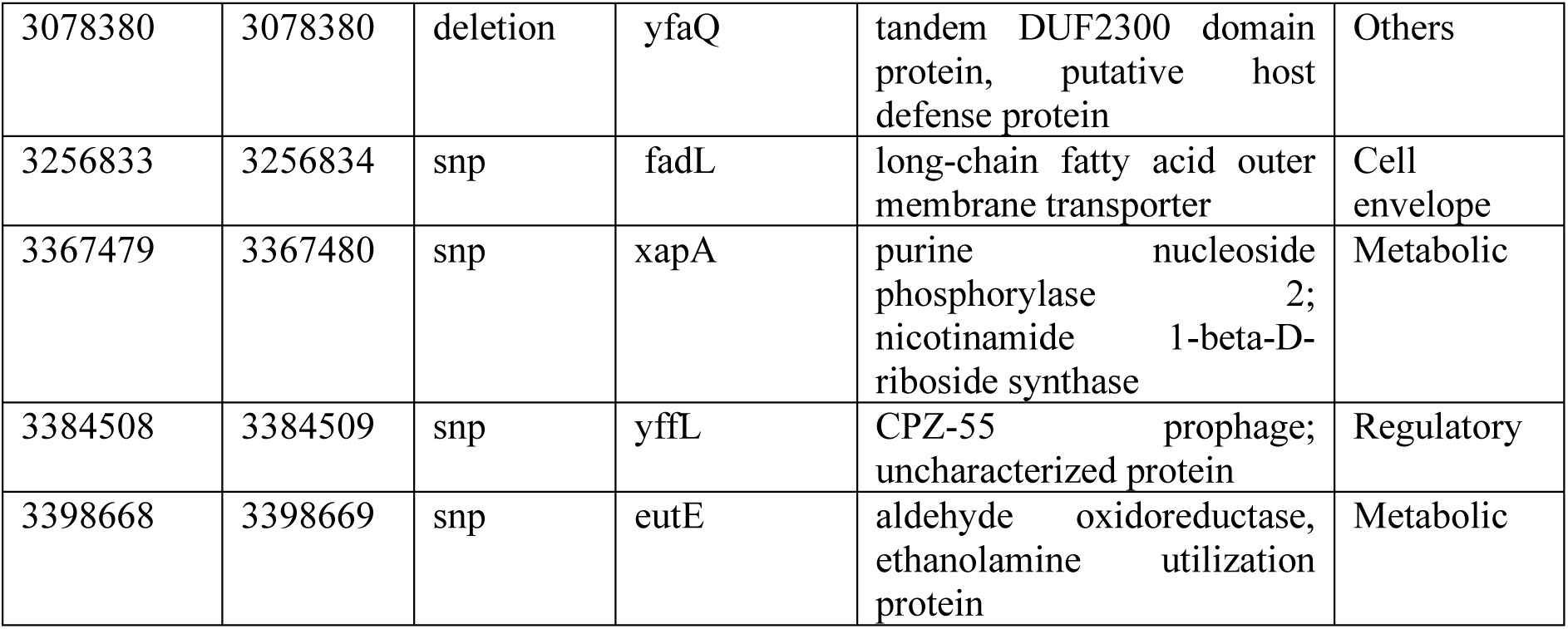
Functional classification of mutations in the Metoclopramide-evolved trajectory Trj1 (folB inhibition).

**Supplementary Table 11.**
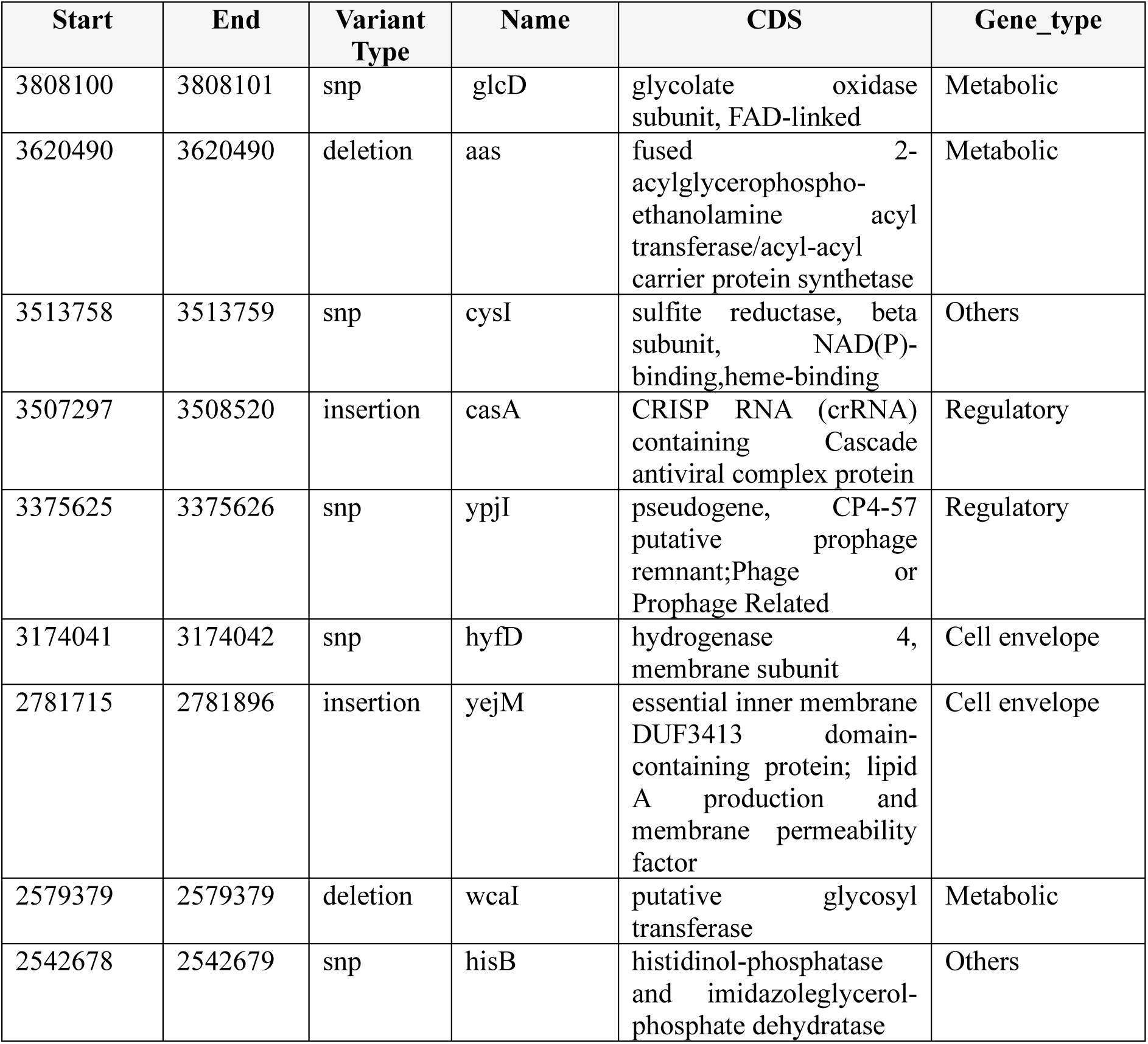
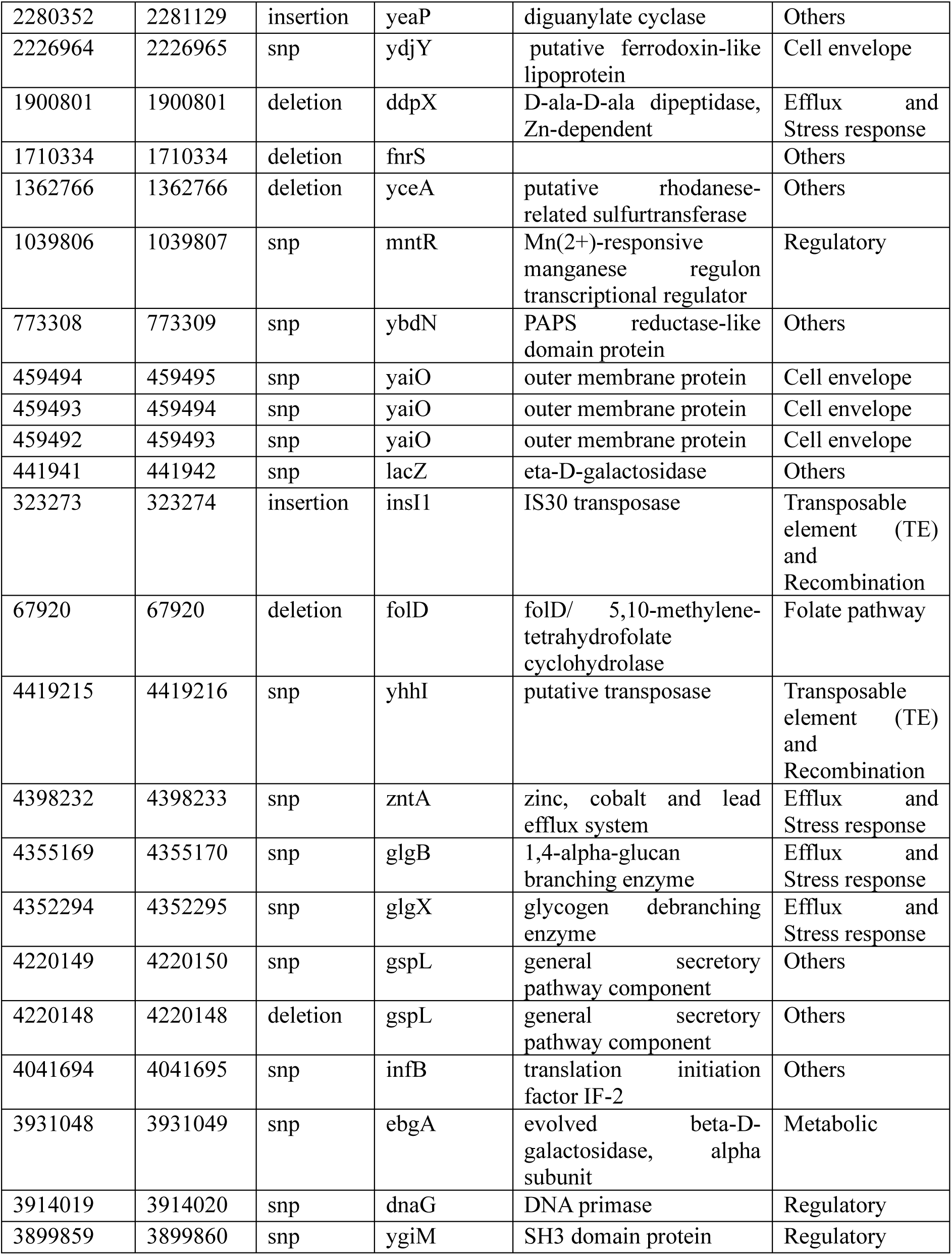
Functional classification of mutations in the Metoclopramide-evolved trajectory Trj2 (folB inhibition).

**Supplementary Table 12.** Functional classification of mutations in the Metoclopramide-evolved trajectory Trj3 (folB inhibition).

| Start | End | Variant Type | Gene Name | CDS | Gene_type |
| --- | --- | --- | --- | --- | --- |
| 3808100 | 3808101 | snp | glcD | glycolate oxidase subunit, FAD-linked | Metabolic |
| 3620490 | 3620490 | deletion | aas | fused 2-acylglycerophospho-ethanolamine acyl transferase/acyl-acyl carrier protein synthetase | Metabolic |
| 3513758 | 3513759 | snp | cysI | sulfite reductase, beta subunit, NAD(P)-binding, heme-binding | Metabolic |
| 3507297 | 3508520 | insertion | recN | recombination and repair protein | Transposable element (TE) and Recombination |
| 3375625 | 3375626 | snp | ypjI | pseudogene, CP4-57 putative prophage remnant | Regulatory |
| 3174041 | 3174042 | snp | hyfD | hydrogenase 4, membrane subunit | Cell envelope |
| 2781715 | 2781896 | insertion | yejM | essential inner membrane DUF3413 domain-containing protein; lipid A ion and membrane permeability factor | Cell envelope |
| 2579379 | 2579379 | deletion | wcaI | putative colanic biosynthesis glycosyl transferase | Metabolic |
| 2542678 | 2542679 | snp | hisB | histidinol-phosphatase and imidazoleglycerol-phosphate dehydratase | Metabolic |
| 2280352 | 2281129 | insertion | yeaP | diguanylate cyclase | Others |
| 2226964 | 2226965 | snp | ydjY | putative ferredoxin-like lipoprotein | Cell envelope |
| 1900801 | 1900801 | deletion | ddpX | D-ala-D-ala dipeptidase, Zn-dependent | Metabolic |
| 1710334 | 1710334 | deletion | dbpA | ATP-dependent RNA helicase, specific for 23S rRNA | Regulatory |
| 1362766 | 1362766 | deletion | yceA | putative rhodanese-related sulfurtransferase | Metabolic |
| 1039806 | 1039807 | snp | mntR | Mn(2+)-responsive manganese regulon transcriptional regulator | Regulatory |
| 773308 | 773309 | snp | ybdM | Spo0J family protein, ParB-like nuclease domain | Others |
| 459494 | 459495 | snp | yaiO | outer membrane protein | Cell envelope |
| 459493 | 459494 | snp | yaiO | outer membrane protein | Cell envelope |
| 459492 | 459493 | snp | yaiO | outer membrane protein | Cell envelope |
| 434952 | 434953 | snp | yaiO | outer membrane protein | Cell envelope |
| 323273 | 323274 | insertion | insI1 | IS30 transposase | Transposable element (TE) and Recombination |
| 67920 | 67920 | deletion | lptD | LPS assembly OM complex LptDE, beta-barrel component | Others |
| 4419215 | 4419216 | snp | prlC | oligopeptidase A | Metabolic |
| 4398232 | 4398233 | snp | zntA | zinc, cobalt and lead efflux system | Efflux and Stress response |
| 4355169 | 4355170 | snp | glgX | glycogen debranching enzyme | Efflux and Stress response |
| 4352294 | 4352295 | snp | glgX | glycogen debranching enzyme | Efflux and Stress response |
| 4220149 | 4220150 | snp | gspL | general secretory pathway component, cryptic | Others |
| 4220148 | 4220148 | deletion | gspL | general secretory pathway component, cryptic | Others |
| 4041694 | 4041695 | snp | infB | translation initiation factor IF-2 | Metabolic |
| 3931048 | 3931049 | snp | ebgA | evolved beta-D-galactosidase, alpha subunit | Metabolic |
| 3914019 | 3914020 | snp | ebgA | evolved beta-D-galactosidase, alpha subunit | Metabolic |
| 3899859 | 3899860 | snp | ygiF | inorganic triphosphatase | Others |

**Supplementary Table 13.** Gene family-level organization of mutations across Metoclopramide-evolved trajectories.

| Gene Family | Trj1 | Trj2 | Trj3 | Similarity Pattern |
| --- | --- | --- | --- | --- |
| glc* | glcB | glcD | glcD | Same metabolic family |
| cys* | cysC | cysI | cysI | Sulfur assimilation family |
| his* | hisS | hisB | hisB | Histidine pathway family |
| yej* | – | yejM | yejM | Same gene |
| ydd/ydj* | – | ydjY | ydjY | Same gene |
| glg* | – | glgB, glgX | glgX | Glycogen metabolism family |
| gsp* | – | gspL | gspL | Secretion system family |
| ins* | — | insI1 | insI1 | Transposable element family |
| mntR / metal regulation family | — | mntR | mntR | Same gene |
| ddp* | — | ddpX | ddpX | Dipeptide metabolism family |
| yaiO / envelope gene | — | yaiO | yaiO | Same gene |
| wca* | — | wcaI | wcaI | Capsule biosynthesis family |

